# Argonaute 15 regulates immunity of potato to filamentous pathogens through transposon-derived small RNAs

**DOI:** 10.64898/2026.09.25.754549

**Authors:** Shailja Singh, Xinyi Hu, Kristian Persson Hodén, Zhen Liao, Christina Dixelius

## Abstract

Canonical ribonucleases such as Argonaute (AGO) participate in RNA-based gene regulatory processes. They are important for small RNA (sRNA) association and the formation of RNA-induced RNA silencing complexes. Potato (*Solanum tuberosum*) has 14 *AGO*-encoding genes, including the *Solanum*-specific *StAGO15*, which is activated upon infection by filamentous pathogens. The resistant phenotype conferred by ectopic expression of *StAGO1*5 was abolished in plants in which a 21-nt sequence in the PAZ-encoding region of *StAGO15* was targeted using an artificial miRNA. We then analyzed the sRNAs associated with *StAGO15* in *Phytophthora infestans* leaf infection. Small RNAs were coimmunoprecipitated, subjected to deep sequencing, and analyzed against the *P. infestans* and potato genomes, in which cleavage sites were predicted by applying the R package, smartPARE. The most notable difference between pathogen-and water-inoculated *StAGO15* samples was observed in the transposon (TE) category. Here, two categories of nonautonomous TE dominated: the miniature inverted-repeat transposable elements (*MITEs*) and short interspersed nuclear elements. The *StAGO15*-associated sRNAs bind and target complementary endogenous potato RNAs. One example is siRNA122, which acts on a cleavage site in the resistance gene PGSC0003DMG400015681 (*StRES1*). siRNA122 is derived from a *MITE*; upon deletion of this source sequence, resistance is restored. Our findings provide new insights into the plant immune surveillance network and demonstrate that deleting a 21 nt RNA sequence from a *MITE* transposon restored a defense response against two important filamentous pathogens.

**Significance Statement:** Potato is a major food crop in the world with increasing importance in the developing countries due to its high adaptive capability to variable climate conditions. The potato plant is known to be attacked by many pathogens and insects during its growing season. Here we demonstrate the importance of a small RNA affecting the defense level to the late and early blight diseases which provides new possibilities for crop improvement.

## INTRODUCTION

Potato (*Solanum tuberosum*) is the largest non-cereal food crop worldwide, ranking fourth in production after rice, wheat and maize (FAOSTAT, 2023). This position can be explained by its wide adaptability to chiefly temperate growing conditions, high yield potentials, and advantageous nutritional composition. Potato tubers serve as the primary nutrient storage and reproductive organs of the plant; therefore, potato breeding goals emphasize tuber quantity and quality unlike the research on cereals and its focus on seed improvement (Bradshaw, 2021). Besides playing an important role in food production, potato tubers have become a valuable industrial raw material for bioplastics, packaging coatings, adhesives, and starch gelatinization among several applications (Ahmad et al., 2024). The first potato genome was published in 2011 by the International Potato Genome Sequencing Consortium (PGSC) after sequencing a doubled monoploid genotype (The Potato Genome Sequencing Consortium, 2011). This was a considerable effort, and the assembly has since undergone several genome revisions (https://spuddb.uga.edu).https://spuddb.uga.edu potato comprises 48 chromosomes (*2n* = *4x* = 48). The estimated genome size is 2.67 Gb with more than 125,077 (99.21%) annotated protein-coding genes (Wang et al., 2022). Based on an extensive analysis of 24 wild and 20 cultivated diploid potatoes, a complex evolutionary pattern has been revealed (Tang et al., 2022). Among several interesting findings, including large chromosomal inversions, was the expanded numbers of disease-resistance genes from 478 in *Solanum morelliforme* to 1,976 in *Solanum chacoense.* Potato is affected by several diseases, among which potato late blight caused by the oomycete pathogen *Phytophthora infestans*, is the most important biotic constraint to potato production worldwide. Great efforts to introduce new resistance genes from related *Solanum* species have been undertaken for the last 100 years but the exceptional adaptation capacity of *P. infestans* constantly results in new virulent strains eroding the introduced resistance traits (Vleeshouwers et al., 2011; Paluchowska et al., 2022). Improving the broad-spectrum durability of late blight resistance is presently an important breeding goal. However, genetic progress is laborious owing to the tetraploid nature of potato, its complex gene families, and high genetic heterozygosity, which has impeded the identification of valuable individual plants (Qu et al., 2024).

Noncoding small RNAs (sRNAs) are components of eukaryotic and prokaryotic gene regulatory processes, which include transposon activities. sRNAs and their effects on gene activity were initially reported in viruses (Hamilton & Baulcombe, 1999; Baulcombe, 2004), and subsequent studies were rapidly conducted in other organisms including plants, particularly *Arabidopsis thaliana*. Many sRNA classes are described today with different lengths, biogenesis, functions, and targets (Axtell, 2013; Zhan & Meyers, 2023). In plants, small RNAs are mainly classified as either microRNAs (miRNAs) or small interfering RNAs (siRNAs) based on differences in their precursors and biogenesis. miRNAs regulate target mRNAs *via* post-transcriptional gene silencing, whereas siRNAs induce transcriptional gene silencing. The outcome of these two silencing pathways is known to regulate developmental processes, control transposon activity, and respond to different stress conditions. This gene regulation is mediated by a few canonical ribonucleases that participate in the initiator and effector steps (Borges & Martienssen, 2015; Axtell & Meyers, 2018), among which Argonautes (AGOs) constitute an important category. AGOs play important roles in sRNA association and the formation of RNA-induced RNA silencing complexes (RISCs). This function is undertaken by different AGO protein domains that bind diverse sRNA classes.

The Arabidopsis genome contains ten AGO proteins divided into three distinct phylogenetic clades based on the amino acid sequences; AGO1/AGO5/AGO10, AGO2/AGO3/AGO7, and AGO4/AGO6/AGO8/AGO9 (Vaucheret, 2008). Broadly, the three clades can be further divided based on their functions; members of the AGO1/5/10 clade are RNA slicers, AGO2/3/7 bind sRNAs, and AGO4/6/8/9 are chromatin modifiers (Montgomery et al., 2008; Havecker et al., 2010; Ji et al., 2011). The functions of AGO proteins are known to be more diverse today than previously believed, as their roles have been attributed to transcriptional and posttranscriptional gene silencing guided by 24-or 21-nt sRNAs (Ma & Zhang, 2018; Liang et al., 2023). In plant defense, AGO4/6/8/9 in Arabidopsis are known to play important roles in response to viruses and bacteria (Martín-Merchán et al., 2023). Systemic acquired resistance to infection by the bacterium *Pseudomonas syringae* requires AGO4, an Argonaut that also functions in the RdDM pathway (Agorio & Vera, 2007; Rebolledo-Prudencio et al., 2022). In contrast, Arabidopsis AGO2 promotes pathogenesis-related protein 1 responses (Zhang et al., 2011), suggesting overlapping functions between different AGO clades in response to different invading pathogens. Extensive genomic analysis has revealed rapidly increasing numbers of AGO proteins, refining their evolution in Viridiplantae and providing new insights into their function (Julien et al., 2020; Li et al., 2022; Bélanger et al., 2023).

The gene families in *Solanaceae* vary in size because of duplications, and the speed and mode of evolutionary events (Wang et al., 2018). This can be exemplified by the *AGO* gene family. To date, 10 *AGOs* in *Nicotiana obtusifolia* and 17 in *N. tabacum* have been identified (Liao *et al*., 2020). The potato genome contains 14 *AGOs*, whereas the closely related tomato (*S. lycopersicum*) has 15 A*GOs* (Xian et al., 2013). In both species, orthologs of *AGO8* and *AGO9* are missing, but are present in *Nicotiana* species, where AGO8 mediates defense against herbivore attacks (Pradhan et al., 2017). The genera *Solanum* and *Nicotiana* diverged *c*. 24 million years ago (Mya), whereas potato and tomato species split more recently, that is *c*. 8 Mya, demonstrating the dynamic genetic processes, which are further reinforced by various breeding technologies (Särkinen et al., 2013; Bombarely et al., 2016). Members of the AGO4 clade vary within eudicots and are presently divided into three subbranches. Solyc03g117600, located in the AGO4-II subclade (Li et al., 2022), is the closest tomato homolog to AGO15 in potato. In the *Solanum tuberosum* (*St*) potato genome, three AGO4 sequences are present, all of which form a separate subclade together with AGO6 and AGO15 (Liao et al., 2020). *StAGO15* is located on chromosome 3 together with *StAGO1b* (Liao et al., 2020). Among the potato AGOs it is most similar to *StAGO4a* (50% identity), consistent with an origin by ancient duplication within the AGO4 clade.

In our previous study, constitutively expressed *StAGO15* was clearly activated by filamentous pathogen challenge, suggesting that *StAGO15* plays an important role in the immune response of potato (Liao et al., 2020). In this study, we infected transgenic potatoes harboring *StAGO15-GFP* with *Phytophthora infestans* (*Pi*) followed by coimmunoprecipitation, sequencing, and analysis of the sRNA-associated population to clarify possible new sRNA functions during potato infection. The datasets were processed in the smartPARE pipeline to predict sRNA classes and cleavage sites (Persson Hodén et al., 2021). TE-derived sRNAs were predominantly predicted to cleave resistance genes. *StAGO15-*overexpressing potato plants showed resistance to both *P. infestans* and *Alternaria solani* compared to the wildtype, whereas the defense responses of *StAGO15*-knockdown plants were compromised. Gel shift assays demonstrated binding between StAGO15 and siRNAs. A CRISPR/Cas9 construct designed to target 21 nt siRNAs located in a *MITE* transposon was assayed transiently in potato, resulting in resistance responses to both filamentous pathogens. The explanations of the findings are discussed and illustrated using a model.

## RESULTS

### Partition of potato and *P. infestans* sRNAs

We infected the leaves of transgenic potato plants harboring *p35S:StAGO15-GFP* with *P. infestans* (strain 3928A) or water and collected the samples in triplicate at 5 days post-inoculation (dpi), a time point corresponding to the biotrophic-necrotrophic transition when *StAGO15* shows peak transcriptional activation (Liao et al., 2020). Coimmunoprecipitated sRNAs were collected and analyzed on an Illumina NextSeq 2000 platform. Computational processing revealed three refined sRNA groups: potato sRNAs from the leaves of *p35S:StAGO15-GFP* plants inoculated with the pathogen or H_2_O, and sRNAs of *P. infestans* from the inoculated transgenic potato leaves (Table S1). The potato size profiles were centered at approximately 22 nt to 24 nt, with a peak of 23 nt in the H_2_O-inoculated samples and a clear enrichment of 22 nt sRNAs in the pathogen-inoculated samples (Fig. 1a). Plant sRNAs range from 18 to 24 nt in size, with a typical predominance of 21 and 24 nt sRNAs (Axtell, 2013; Chávez Montes et al., 2014). Canonical miRNAs are 20 to 22 nt in length (Ghildiyal & Zamore, 2009). Earlier, the 21 nt size class in potato was found to dominate the interaction with a *P. infestans* strain harboring *PiAgo1-GFP* (Hu et al., 2022). When monitoring the distribution of PisRNAs in the StAGO15 material, bimodal peaks were identified, one at 21–22 nt and the other at 25–26 nt (Fig. 1b). Peaks of different sizes, 18, 21, and 24 nt, during potato infection with *PiAgo1-GFP* were previously observed (Hu et al., 2022). Recognition of the 5′ terminal nt is known to form the basis for AGO sRNA specificity in Arabidopsis and other model organisms (Mi et al., 2008; Montgomery et al., 2008; Wu et al., 2020). In this study, *StAGO15* presented a high and stable level of 5′ A (adenine) during infection, whereas a slightly increased percentage of 5′G (guanine) and a decrease in 5′U (uracil) compared with those in the control condition were observed (Fig. 1c). In the alignment of StAGO15 with AGO1 from Arabidopsis, sequence divergences were observed in the PIWI domain (Liao et al., 2020). Notably, the catalytic “slicing” tetrad D-E-D-H/D was changed to a G-E-Q-R motif. Furthermore, the nucleotide specificity loop (NSL) differed from that of AtAGO1, and these differences may jointly contribute to the specific function(s) of StAGO15. Such structural and sequence divergence is particularly significant, as small PIWI domain variations can tune small RNA targeting and AGO proteins can exhibit diverse functions beyond canonical slicing, including altered substrate specificity (Wu et al., 2015; Xiao et al., 2023), highlighting the potential for StAGO15’s divergent PIWI domain to mediate distinct molecular mechanisms. *AtAGO5* is the only plant AGO reported so far with a less strict 5′ terminal association of nucleotides, which varies between C (cytosine), A, and G (Frank et al., 2012). How pathogen-induced stress influences the choice of 5′ preference in the host remains unclear. In *P. infestans,* different AGOs have different 5′ preferences (Åsman et al., 2016), and *PiAgo1* shifts the 5′nt preference from C to U during the infection of potato (Hu et al., 2022). The enrichment of the U 5′nt preference of *PisRNAs* observed here (Fig. 1c) is in line with earlier reports (Vetukuri et al., 2012; Åsman et al., 2016). The 5′ specificity of *AtAGO4* and *AtAGO6* is determined by the coordination of the MID and PIWI domains and not by the nucleotide specificity loop in the MID domain (Liu et al., 2022). We can only speculate that the somewhat altered folding of the StAGO15 protein and the hydrophobic Phe in the nucleotide specificity loop (Liao et al., 2020) lead to conformational changes that influence the sRNA binding properties, thereby the sRNA sorting.

**Fig. 1.**
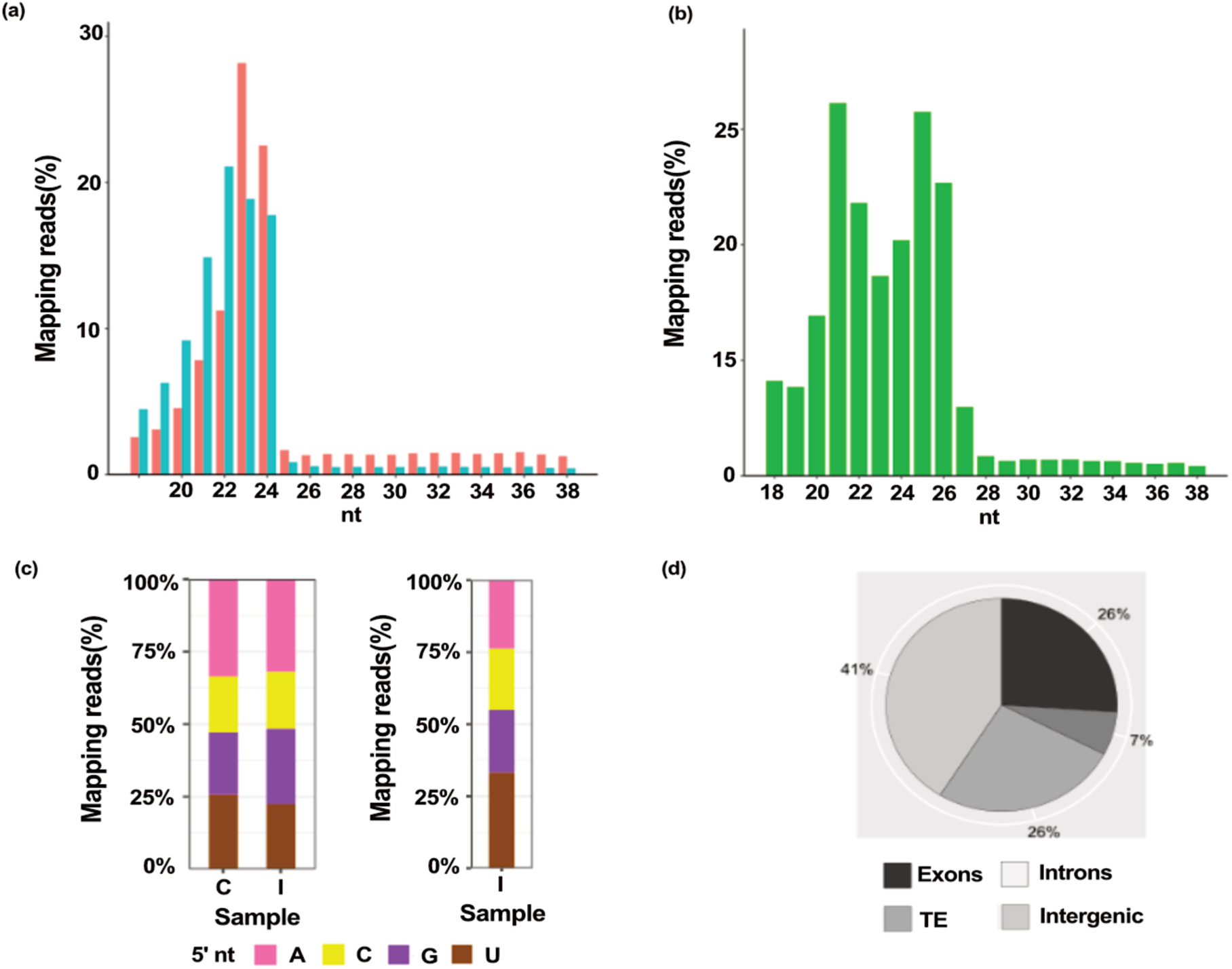
Characteristics of small RNAs derived from coimmunoprecipitated and sequenced RNA samples of potato and *Solanum tuberosum* (*St*) *StAGO15-GFP* leaves. Samples were collected at 5 days post-inoculation (dpi) with *Phytophthora infestans (Pi*). (**a**) Distribution of *St*-sRNAs upon inoculation of *P. infestans* (blue, 11437889 reads) and H_2_O (red, 13202959 reads). (**b**) Distribution of *Pi*-sRNAs (green, 2262964 reads) in inoculated potato. (**c**) Identity of the 5′ terminal nucleotides (nts). Left, *St-*sRNAs. Distribution of the 5′ terminal nucleotide in H2O-inoculated (C) leaves: A: 34%, C: 19%, G: 21%, U: 26%. Distribution of the 5′ terminal nucleotide in *P. infestans-*inoculated (I) leaves: A: 33%, C: 19%, G: 26%, 5′ U: 22%. Right, identity of the 5′ terminal nucleotides of the *Pi-*sRNAs at 5 dpi: A: 24%, C: 21%, G: 21%, U: 34%. Some sRNAs, starting with specific nucleotides, are used to measure specific nucleotide ratios. (d) *St*-sRNAs are derived from the potato genome (Hardigan et al., 2016). The alignment distribution is as follows: 321,161 sRNAs to transposable elements (TEs), where retrotransposons dominate (32% of all TEs), 502,444 sRNAs to intergenic regions, and 399,253 sRNAs to genes, with a distribution of 318,885 sRNAs to exons and 80,368 sRNAs to introns.

### *MITEs* are an essential TE source of potato sRNAs associated with StAGO15

The different potato sRNAs obtained *via* StAGO15 pulldown were derived from intergenic regions (41%), exons (26%), transposons (26%), and introns (7%) (Fig. 1d). Twelve miRNAs were detected in the pathogen-inoculated samples, whereas 20 were identified in the H_2_O-inoculated control samples *via* searches in the miRbase database (Table S2). In an earlier work, we compiled sRNA and degradome-based data from transgenic *StAGO1a* plants, *P. infestans* infected materials including a *PiAgo1-GFP* strain, and wildtype strains in various combinations, to design an R package (smartPARE) for identifying the true cleavage sites associated with this plant‒pathogen interaction (Persson Hodén et al., 2021). The sRNA sequence data generated in the present study from *StAGO15-GFP* transgenic potato challenged with either water or *P. infestans* were integrated into this smartPARE pipeline. The most notable difference between the pathogen-and water-inoculated *StAGO15* samples was observed within the transposon (TE) category (Fig. S1), wherein 100 sRNAs with a TE origin but no phasiRNAs were activated upon infection. The different potato-TE sRNAs obtained *via StAGO15* pulldown originated from two nonautonomous TEs: miniature inverted-repeat transposable elements (*MITEs*; ∼ 80%) and short interspersed nuclear elements (*SINEs*; ∼ 9%) (Table S3). The size distribution of TE-generated sRNAs varied between 19 and 24 nt, with a clear peak of 22 nt in our materials (Table S4). This is a wider size interval than the earlier sequenced Solanaceae gel fragments, which were mainly 24 nt (Kuang et al., 2009).

Next, the targeted potato sequence genes of predicted sRNAs with different normalized fragment abundance (NFA) levels upon infection in comparison with those in the non-infected controls were organized according to the annotated gene functional categories in the potato genome (Fig. S2). More resistance genes were predicted as targets during infection than in the control samples. When the different datasets were compared, more genes were found with increased numbers of sRNAs in the infected samples, potentially leading to an increased number of downregulated mRNAs upon infection (Fig. S3). Bioinformatics analysis of the StAGO15 pulldown data did not reveal any targets of *Pi*-sRNAs in *P. infestans* or potato. Likewise, no potato sRNA was predicted to have targets in *P. infestans.* Taken together, these data suggest that *StAGO15*-associated sRNAs bind and target complementary endogenous potato RNAs.

### Resistance genes can be targeted by TE-derived sRNAs

*StAGO15*-overexpressing (OE) and *StAGO15*-knockdown (KD) plants were used for functional comparison. When scored for pathogen responses, the OE lines presented a clear resistance phenotype, whereas the knockdown lines were susceptible to both *P. infestans* and *A. solani* pathogens (Fig. 2a, Fig. S4). The leaf phenotypes were also supported by pathogen DNA quantification (Fig. 2b) and relative transcript level data (Fig. 2c). Three independent lines expressing *StAGO15* from its native promoter also showed reduced *P. infestans* DNA content relative to the wildtype (Fig. S10). This supports the observation that the phenotype is reproducible under native-promoter-driven *StAGO15* expression and is not restricted to constitutive 35S-driven expression. The latter included KD transgenic plants in which sequences corresponding to the PAZ region were targeted by applying an artificial miRNA approach (Fig. 2d).

**Fig. 2.**
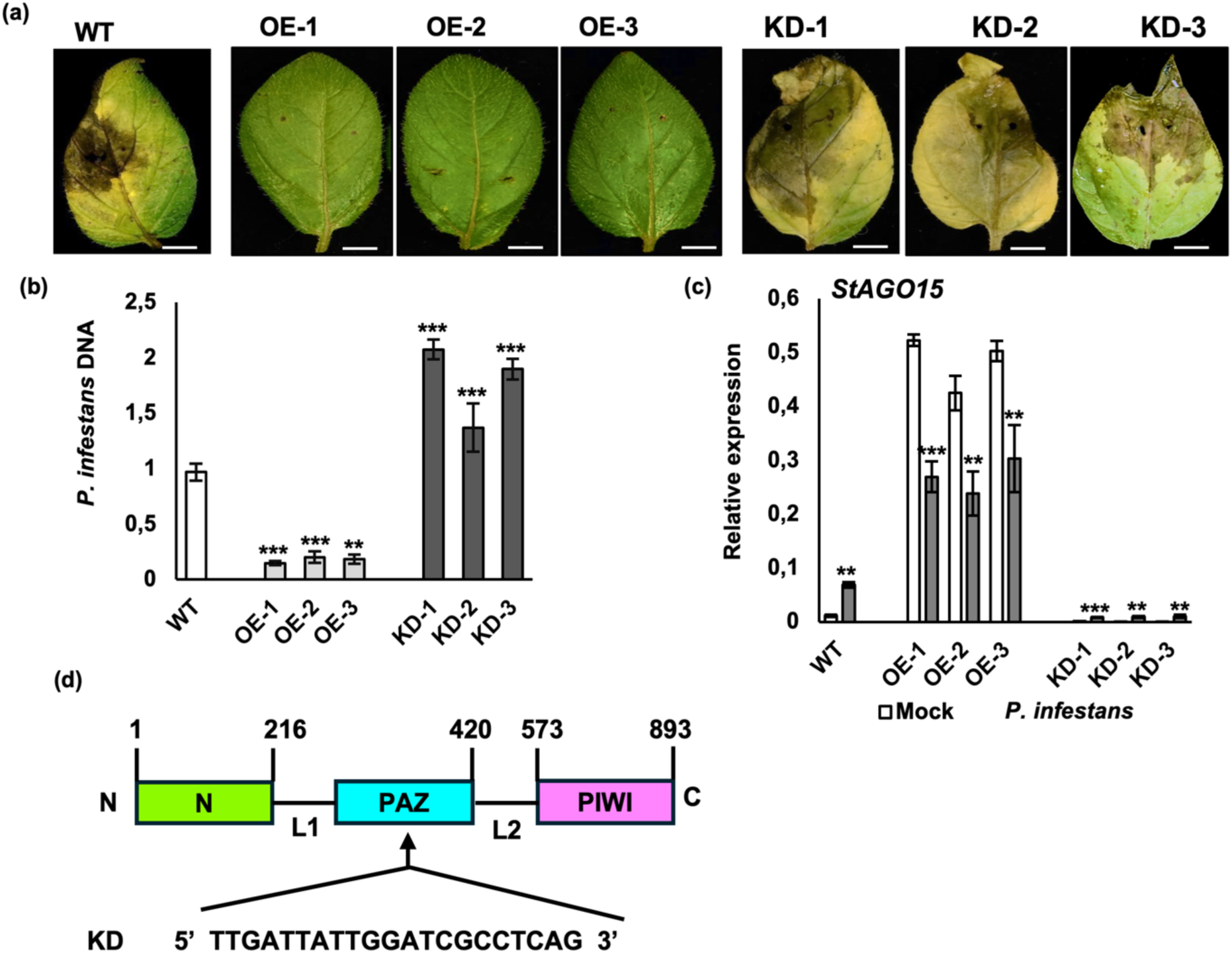
*StAGO15* is essential for potato defense against *Phytophthora infestans*. (**a**) Disease phenotypes of the wildtype (WT; cv. Desirée) and three transgenic overexpression lines containing the *p35S:StAGO15-GFP* construct (OE1, OE2, OE3) and three transgenic knockdown lines containing the *p35S:amiStAGO15-GFP* construct (KD1, KD2, KD3) inoculated with *P. infestans*. Bars, 1 cm. (**b**) Relative *P. infestans* DNA content in leaves from overexpression and knockdown potato plants. *P. infestans* DNA was quantified with *PiO8* primers and normalized to potato *StActin101*. (**c**) Relative expression of *StAGO15* in leaves from the overexpression and knockdown plants inoculated with H_2_O (white) or *P. infestans* (gray). *StActin101* was used as the reference gene. Error bars indicate the mean ± s.d. Asterisks indicate statistically significant differences at *\*P < 0.05, **P < 0.01* and *\*\*\*P < 0.001* according to two-tailed Student’s t tests. n ≥ 3. (**d**) Schematic illustration of the sequence in *StAGO15* targeted by the KD construct and the corresponding domains. N-terminal, Linker 1, PAZ (Piwi/Argonaute/Zwille), Linker 2, C-terminal PIWI (P-element-induced Wimpy).

The next step involved validation of the predicted sRNA targets in potato (Dataset 1). The candidate targets with the highest normalized sRNA abundance values (>1.5 RPM) were selected for RT-qPCR analysis. Thirteen predicted targets presented contrasting expression patterns between wildtype and *StAGO15*-overexpressing potato lines (Fig. S5a, Table S5), and in quantification of the corresponding sRNAs. We were able to generate data for ten siRNAs most of which were more highly expressed in the OE plants compared with those in the wildtype plants (Fig. S5b), whereas the predicted target genes in OE plants in most cases presented decreased expression (Fig. S5a).

Four siRNA candidates: siRNA122, siRNA109, siRNA111, and siRNA118 were selected for detailed downstream analysis based on the following criteria: high normalized enrichment in StAGO15 pulldown libraries under *P. infestans* infection (Dataset 1), strong sequence complementarity with the predicted cleavage sites, favorable unpaired energy (UPE), and low expectation scores (Dataset 2), indicating high target site accessibility and binding affinity. Together, these parameters support their potential as biologically relevant siRNAs. Quantification of these siRNAs in OE plants compared to those in the wildtype also demonstrated their elevated accumulation in OE lines (Fig. S5b). Further, to ensure that *StAGO15* binds siRNAs, mobility shift assays were performed. As expected, binding of the selected siRNA candidates was observed (Fig. S6). Based on the most consistent outcome of the criteria mentioned above, including infection-specific cleavage, siRNA122 was chosen for further analysis.

The predicted siRNA122 cleavage site in the resistance (*R*) gene PGSC0003DMG400015681 (*StRES1*) was validated *via* 5′RACE analysis using total RNA extracted from the leaves of OE plants at 5 dpi with *P. infestans*. The amplified PCR products, ranging from approximately 150-200 bp, were cloned, sequenced, and mapped to the potato genome ver. 6.1 (Pham et al., 2020). Three of twenty sequenced clones terminated at the predicted cleavage site in the candidate *R* gene (Fig. 3a). Similar data were generated *via* 5′ RACE from the other selected siRNA candidates (Fig. S7). The 5′ RACE data were followed by northern hybridization using 21 nt siRNA as a probe, which revealed enrichment of fragments corresponding to this size in OE plants inoculated with *P. infestans* in contrast to those in the KD lines (Fig. 3b). Next the transcript levels of *StRES1* and siRNA122 were compared in OE and KD plants. At 5 dpi, siRNA122 levels were higher in OE plants, whereas *StRES1* transcript levels were lower; in KD plants, siRNA122 levels were lower and *StRES1* transcript levels were higher (Fig. 3c, d). This pattern suggests that siRNA122 may contribute to *StRES1* repression in a StAGO15-dependent manner. Similar changes were observed for two other experimentally validated siRNA-target pairs: siRNA109 and siRNA111 were reduced in OE plants, whereas their resistance-gene targets, PGSC0003DMG402020581 (*StRES2*) and PGSC0003DMG400021887 (*StRES3*), respectively, were elevated (Fig. S5). Together, these observations suggest that altered *StAGO15* expression affects multiple siRNA-target interactions, which may collectively contribute to the observed changes in resistance-gene expression.

**Fig. 3.**
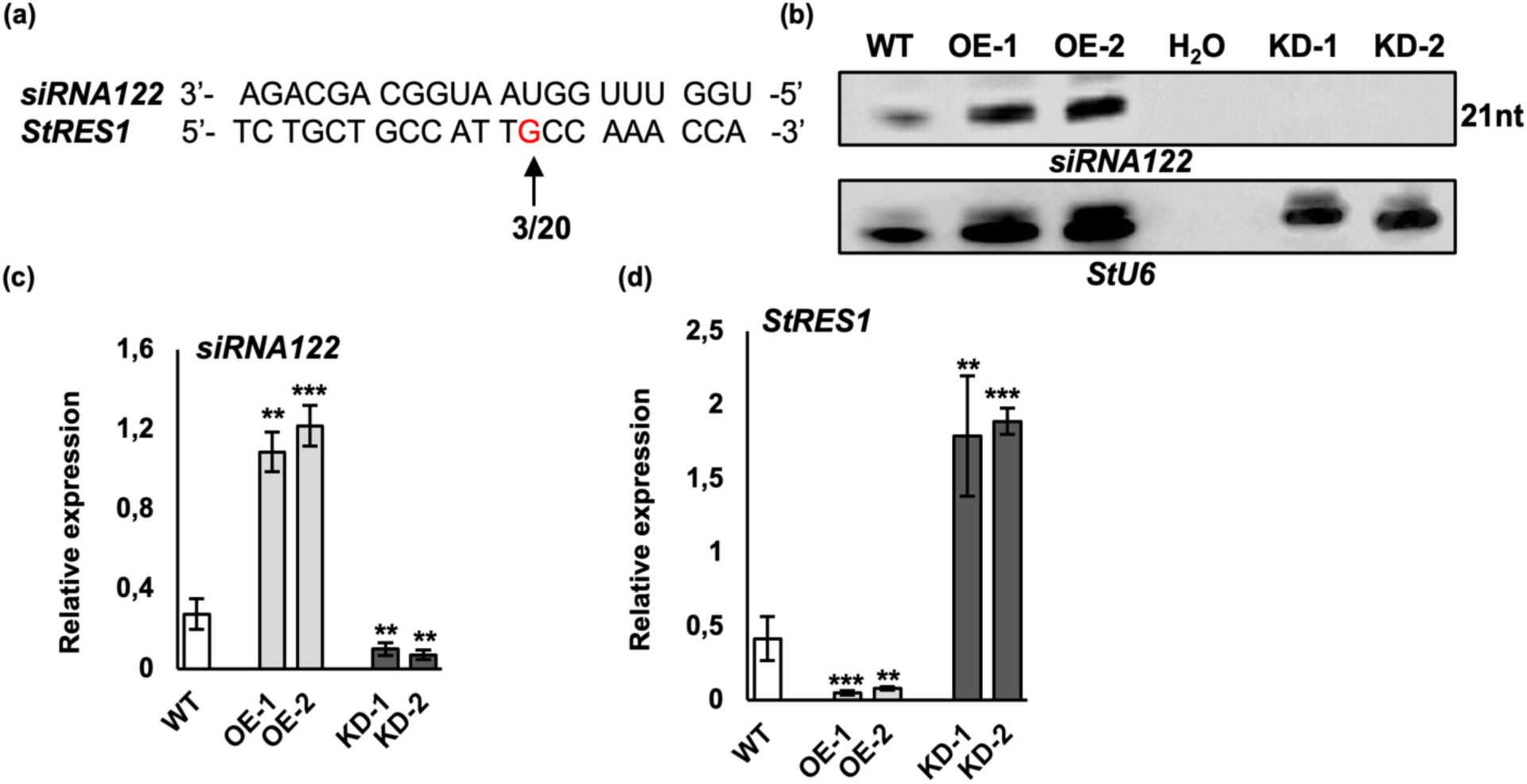
*MITE*-generated siRNA target resistance genes. (**a**) Schematic drawing showing the alignment of siRNA122 with PGSC0003DMG400015681 (*StRES1*) at the predicted cleavage site. The arrow indicates the cleavage efficiency identified using 5′ RACE at the predicted cleavage site upon inoculation of *Phytophthora infestans* and water. Red nt = sequence divergence between cv. Desirée and the reference genome. (**b**) Northern blot analysis using a DIG-labeled RNA probe for 21 nt siRNA. The following RNA was extracted from infected potato leaves collected at 5 dpi: wildtype (WT; cv. Desirée), two overexpression (OE1, OE2, *p35S:StAGO15-GFP*), and two knockdown (KD1, KD2, *p35S:amiStAGO15-GFP*) lines. The lower panel depicts the U6 snRNA from potato (loading control, probe that cross-reacted with potato U6). (**c**) Relative expression of siRNA122 in leaves from the overexpression and knockdown lines compared with that in the wildtype. (**d**) Relative expression of PGSC0003DMG400015681 (*StRES1*) in leaves from the overexpression and knockdown lines compared with that in the wildtype. *StActin101* was used as a reference gene. Error bars indicate the mean ± s.d. (n = 3). Asterisks indicate statistically significant differences at *\*P < 0.05, **P < 0.01* and *\*\*\*P < 0.001* according to two-tailed Student’s t tests.

### *MITE* sequence editing generates enhanced resistance in potato

Nucleotide-binding leucine-rich repeat (NLR) or resistance (R) proteins are intracellular immune receptors that sense effectors delivered by pathogens. Several NLRs in Solanaceae collaborate with a helper protein named NRC (NLR-required for cell death) to restrict plant invasion by pathogens (Wu et al., 2017). We searched for the helper gene for *StRES1* (PGSC0003DMG400015681) and identified *StNRC3a.1* (Sotub05g007670). First, we checked the transcription levels of *StNRC3a.1*, a sequence not targeted by siRNA122 in contrast to PGSC0003DMG400015681 (*StRES1*). However, we were interested to see if this helper gene and its protein were affected if the *R*-gene partner was down-regulated. Compared with control plants, the transcript level of *StNRC3a.1* in the OE plants was lower, whereas the reverse was observed in KD plants (Fig. S8) suggesting that changes in *StAGO15* affect its ability to bind siRNA122, and thereby alter the function of *StRES1.* As a sensor, *StRES1* could influence the expression of *StNRC3a.1*, explaining the expression patterns of *StNRC3a.1* observed in OE and KD plants. Next, we predicted protein‒protein interactions between the NLR protein StRES1 and the predicted helper protein StNRC3a.1 (Figs. S9a and b). Docking analysis revealed a strong interaction between the StRES1 and StNRC3a.1 proteins, with high predicted interaction, reinforcing their potential role in immune signaling. A 2D plot (LigPlot) analysis revealed tryptophan-321 and glycine-323 within the active interaction site (Fig. S9c), suggesting that these residues constitute the main components of StRES1–StNRC3a.1 binding and are most likely critical for enhanced immune signaling upon pathogen infection.

Transient assays in plants are typically performed with *Nicotiana benthamiana*. However, an efficient transient assay is required when examining species-specific genes. In our case, the *N. benthamiana* genome does not harbor the defense genes predicted to be siRNA targets. Therefore, we developed a transient assay using potato leaves to quickly validate primer design. Improved assay consistency was achieved by adjusting the protocol for use in four-week-old potato plants and a 24-h dark treatment post-infiltration, followed by pathogen inoculation. To test the contribution of siRNA122 directly, we used paired sgRNAs to delete a 201 nt region of chromosome 11 (9,393,316–9,393,516) containing the 21 nt siRNA122 source sequence within a *MITE*. Removing the source sequence eliminates siRNA122 production, so the *StRES1* transcript is no longer targeted for cleavage. The constructs were introduced transiently into potato leaves, with empty vector as control. Responses to the two pathogens were monitored daily. A notable decrease in pathogen establishment and growth was observed on the leaves (Fig. 4a, Fig. 4b). Quantification of *StRES1* expression following infection with *P. infestans* and *A. solani* revealed significantly elevated levels in samples from the genome-edited leaves (Fig. 4c). We interpret these data to indicate that loss of siRNA122 gene targeting allows increased *StRES1* expression. Transcript profiling revealed that *StNRC3a.1* was also upregulated in samples from genome-edited potato leaves in response to pathogen challenge, suggesting a coordinated immune response between *StRES1* and *StNRC3a.1* (Fig. 4d).

**Fig. 4.**
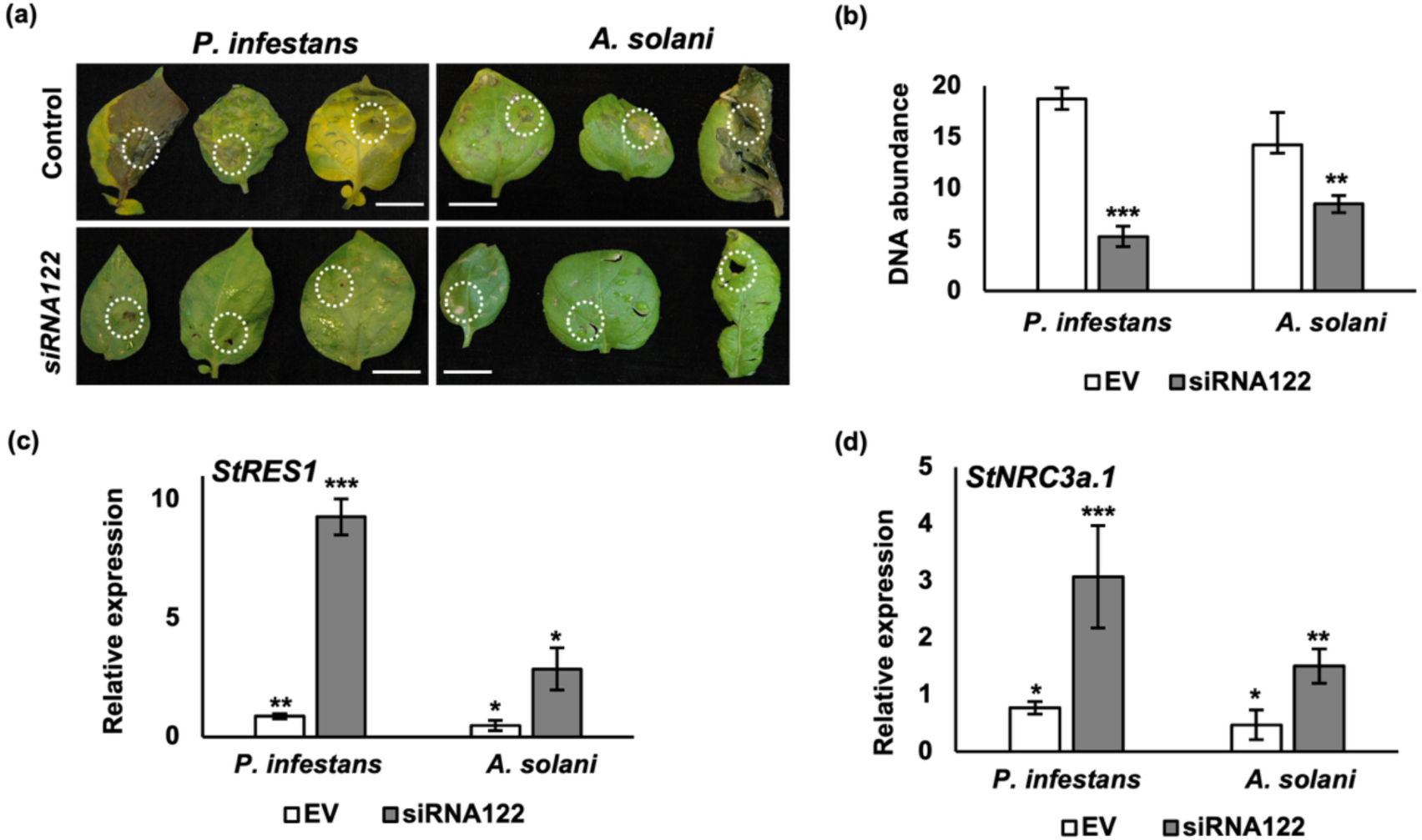
Transient assay in potato, where the siRNA122 targeting *StRES1* is deleted from a *MITE* sequence *via* genome editing. Control plants (empty vector) were challenged with *P. infestans* (left) and *A. solani* (right), with images taken at 5 days post-inoculation (dpi) and 8 dpi, respectively. The dotted circles indicate the inoculation sites. (**a**) Phenotypes of genome-edited plants challenged with *P. infestans* and *A. solani*. Scale bars = 5 mm. (**b**) Relative pathogen DNA quantification in control and genome-edited siRNA122 leaves. *P. infestans* and *A. solani* DNA were quantified with *PiO8* primers and the *A. solani*-specific ITS primers *AsFor* and *AsRev* (Leiminger et al., 2015), respectively, and normalized to potato *StActin101*. Relative expression of (**c**) *StRES1* and (**d**) *StNRC3a.1* in samples from control and genome-edited siRNA122 leaves. *StActin101* was used as a reference gene. Error bars indicate the mean ± s.d. (n ≥3). Asterisks indicate statistically significant differences at *\*P < 0.05, **P < 0.01* and *\*\*\*P < 0.001* using two-tailed Student’s *t*-test.

## DISCUSSION

In this work we followed up on the earlier studies related to StAGO15 in potato (Liao et al., 2020). When *StAGO15* is ectopically expressed in potato, a shift from susceptibility to resistance against filamentous pathogens was observed. An elevated resistance response was also seen when the native promoter was used; however, coimmunoprecipitation using these potato lines did not generate sRNAs that passed the quality control for the sequencing step. Nevertheless, expanding the *StAGO15* gene family or boosting gene activity resulted in pleiotropic defense effects. To follow up the sequencing data, the OE plants were used in the continued analysis.

The different sRNA groups obtained *via* StAGO15 pulldown and sequencing were derived from intergenic regions (41%), exons (26%), transposons (26%), and introns (7%) (Fig. 1d). In the transposon group, two nonautonomous TE-categories dominated: the miniature inverted-repeat transposable elements (*MITE*s; 80%) and short interspersed nuclear elements (*SINE*s; 9%). The most notable difference between pathogen-and water-inoculated *StAGO15* samples was observed within the transposon (TE) category. This led us to closely examine *MITE*s and *SINE*s and their potential regulatory effects on defense responses.

Species in the Solanaceae family have diverse evolutionary histories in which genome duplication and the amplification of genes and transposons play major roles in shaping the genomes (Sanatombi, 2024). In an analysis of 296 *Solanum* accessions, the TE number and composition were found to vary based on the origin and propagation history of different ancestors (Bozan et al., 2023). The TE content was estimated to be 75.5%, with a bias towards retrotransposons, a ubiquitous class of TEs in plants. In terms of copy numbers in the genomes, two groups stand out: abundant long terminal repeat (LTR) retrotransposons and *MITE*s, which are derivatives of autonomous terminal inverted repeat TEs (Pulido & Casacuberta, 2023). TE classes are asymmetrically distributed in the potato genome. LTR retrotransposons are clustered in pericentromeric regions, whereas *MITE*s are mainly located in gene-rich subtelomeric regions (Zavallo et al., 2020). More than 50% of non-LTR retrotransposons and DNA-based TEs are within or close to the nearest genes, suggesting a high potential to impact transcription if activated. Regarding *StAGO15*, *MITE*s are abundant in its vicinity on the potato chromosome 3 but are also frequently distributed on chromosome 11, where *StRES1* is located together with the *MITE* generating siRNA122.

To reduce potential genome rearrangement and mutation frequencies, various systems work to maintain genome stability and minimize gene disruption (Deniz et al., 2019). At least four routes are known in plants to control TE transposition involving methylation processes as well as posttranscriptional and transcriptional gene silencing (Mhiri et al., 2022). The RNA-directed DNA methylation (RdDM) pathway plays a central role in these TE-silencing actions in addition to orchestrating potential gene disruptions (Erdmann & Picard, 2020). However, information on the biological function(s) of RdDMs in potato remains limited beyond the impact of DNA methylation related to abiotic stress (Sabbah et al., 1995; Dutta et al., 2022), tuberization (Ai et al., 2021), and skin phenotype (Kumar et al., 2023).

In the nightshade family, sRNA classes, like the overall genome, have experienced complex evolutionary events, which at least partly explain the present shared and lineage-specific dynamic landscape of sRNAs (Baldrich et al., 2022). The enriched *PHAS* loci in tomato and potato genomes are primarily generated from the abundant resistance genes encoding nucleotide-binding domains and leucine-rich repeats. Expression of the vast number of *R* genes in plant genomes is hypothesized to be controlled by TEs, and epigenetic mechanisms such as methylation can be used to avoid the negative side effects of a constitutively resistant plant (Cambiagno et al., 2021). Potato has a low abundance of 24 nt siRNAs (Persson Hodén et al., 2021; Baldrich et al., 2022). This finding was attributed to the long-term asexual propagation circumventing meiosis, resulting in potential depletion of heterochromatic siRNAs (hc-siRNAs) in the genome. This contrasts with sexually propagated species such as Arabidopsis, which exhibit canonical RNA-directed DNA methylation, in which 24 nt siRNAs are formed by DICER-LIKE 3, bound by AGO4, and further processed to methylate DNA (Papareddy et al., 2020). The *MITE*-derived sRNAs recovered here ranged from 19 to 24 nt with a peak at 22 nt (Table S4), consistent with the sRNA pool available in this species rather than with the 24 nt class that drives RdDM in Arabidopsis. The RdDM pathway in potato is still not fully understood; the same is true of stress-related triggers and their target events. Furthermore, numerous miRNA clusters are common across Solanaceae genomes. A *MITE*-derived miRNA-like siRNA was found for the Solanaceae-specific miR5303 cluster and isomiRs with numerous predicted genome targets (Baldrich et al., 2022). How sRNAs embedded in TE sequences respond in different situations (developmental stages, stresses, and other changes) is unclear and adds to the exceedingly diverse processes together with sRNAs derived from the remaining genomic parts (Vaucheret & Voinnet, 2024). In addition, TEs such as *MITE*s may affect gene functions on their own (Kuang et al., 2009).

We believe that the susceptibility of the wildtype (cv. Desirée) is attributed to siRNA-derived *R*-gene disruptions triggered by biotic stress. A gene-targeting process that could be promoted by the actions of several sRNAs may lead to increased susceptibility (Fig. 5a). Manipulated *AGO* sequences may destabilize the RNA-induced silencing complex (RISC), in turn impacting activity, loading competition, turnover rates, or RNA complementarity and thereby changing phenotypic responses compared with those of the wildtype plants (Li et al., 2019; Iwakawa & Tomari, 2022; Xu & Chen, 2023). In our case, we may also have altered StAGO15 specificity, generating off-target actions that contribute to the resistant phenotype in OE plants, whereas nonfunctional StAGO15 promotes disease development. In the OE plants, *StRES1* remained cleaved while siRNA109 and siRNA111 were reduced and their resistance-gene targets PGSC0003DMG402020581 (*StRES2*) and PGSC0003DMG400021887 (*StRES3*) were elevated (Fig. S5), indicating that the resistant phenotype is not carried by *StRES1* alone but by the distribution of silencing across several targets (Fig. 5b). In the KD plants, where siRNA loading specificity is impaired, this distribution is inverted; *StRES1* accumulates while the plants remain susceptible (Fig. 5c). To this end, the novel PIWI domain of StAGO15 may interfere with the sRNA sorting and redirect sRNAs to other AGOs as reported in Arabidopsis (Zhang et al., 2014). Searching for transcription factor motifs in the 3 kb promoter sequence of *StAGO15* did not reveal any obvious binding site that could possibly participate in this stress response. The transient deletion of the siRNA122 source sequence differs in scope from these transgenic lines, since it removes one *MITE*-derived siRNA while leaving the silencing machinery intact (Fig. 5d).

NLR-mediated immunity is a variable system based on domain effector recognition and activation *via* multiple combinations in a flexible fashion (Ahn et al., 2023; Shepherd et al., 2023). How the endogenous sRNA system has evolved to interfere with this dynamic defense system remains to be clarified. Clearly, interfering with a sequence corresponding to the NLR-NRC protein binding sites would be more cost-effective than targeting two individual genes. In our case, siRNA122 targets a region in *StRES1* corresponding to an NB-ARC and P-domain overlapping sequence (Fig. 5e; Table S6), a common resistance gene target enforced by several miRNAs (López-Márquez et al., 2023). In KD plants, where *StAGO15* transcription is reduced, direct siRNA-mediated suppression of *StNRC3a.1* is less likely owing to fewer available StAGO15-siRNA complexes. This would lead to the activation of *StNRC3a.1*, as its suppression by siRNAs is diminished, a scenario discussed by Liang et al. (2023). Much remains to be clarified regarding the function of a helper-sensor gene pair in the context of sRNAs as well as in the overall context of its regulation, networks, and associated defense signaling. Taken together, our study demonstrates that modifying specific *StAGO15* or *MITE* sequences could create valuable traits to counteract the rapid breakdown of introduced resistance genes in potato, which is of particular importance in the context of *P. infestans*.

**Fig. 5.**
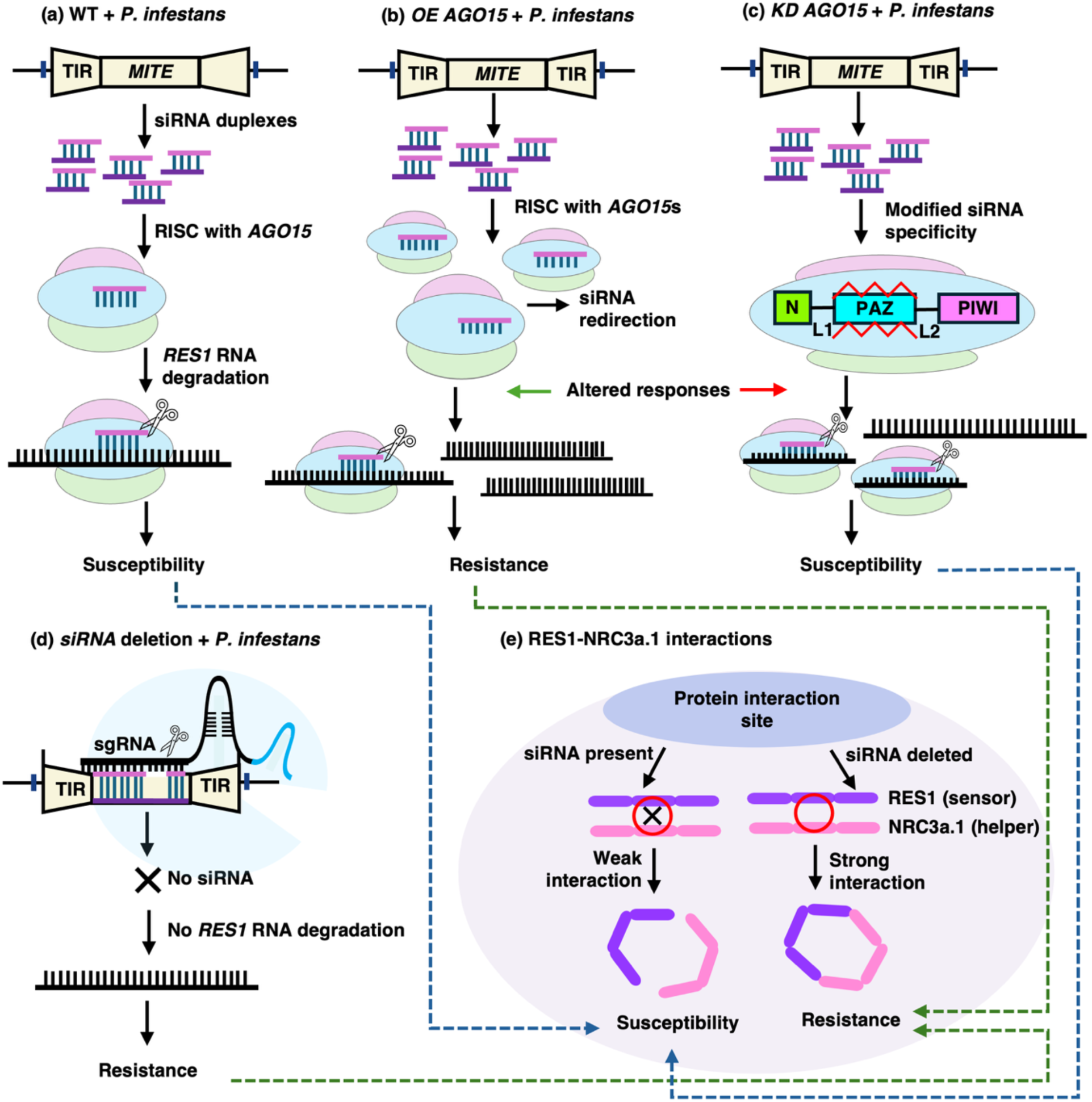
Summary of StAGO15 studies: a hypothetical model. Miniature inverted-repeat transposable elements (*MITE*s) are truncated derivatives of autonomous DNA transposons containing terminal inverted repeats (TIRs) flanked by short direct repeats. Panels (**a-e**) illustrate the proposed mechanistic outcomes under different genetic or experimental conditions involving *MITE*-derived siRNA122, AGO15, and the immune components StRES1 and StNRC3a.1 in potato. (**a**) Wildtype (WT) scenario: Upon challenge with *P. infestans*, siRNA duplexes from the 9393316-9393516 region on potato chromosome 11 are released, and siRNA122 is incorporated into the RISC complex together with AGO15. The complex targets the *StRES1* transcript (PGSC0003DMG400015681), which encodes a TIR-NB-LRR resistance protein. Cleavage occurs 961 bp downstream of the start codon, affecting the NB-ARC domain, and the transcript is degraded, resulting in a susceptible phenotype. (**b**) Ectopic AGO15 expression (OE + *P. infestans*): Increased AGO15 abundance promotes the loading of *MITE*-derived siRNA duplexes into RISC. *StRES1* is cleaved, whereas other siRNAs derived from the other *MITE* regions are redirected away from their targets simultaneously, allowing accumulation of the StRES2 and StRES3 transcripts, which leads to a resistant phenotype. (**c**) *StAGO15* knockdown (KD + *P. infestans*): A 21 nt deletion in the PAZ-encoding region encoded by *StAGO15* causes a frameshift mutation and premature stop codon, together resulting in a truncated, non-functional AGO15 protein in the KD plants. As the PAZ domain is essential for anchoring and orienting siRNAs during the RISC assembly, its loss impairs the specificity of siRNA loading. The distribution of silencing across the three targets is inverted relative to OE; *StRES1* escapes cleavage and accumulates, whereas *StRES2* and *StRES3* are targeted and degraded, thereby resulting in a susceptible phenotype. (**d)** siRNA122 deletion (+ *P. infestans*): CRISPR-Cas9-mediated deletion of the 20 bp *MITE*-derived siRNA122 (TCTGCTGCCATTACCAAACCA) sequence eliminates siRNA122 production. Therefore, *StRES1* mRNA is no longer targeted or downregulated, which leads to restored resistance. **(e)** StRES1-StNRC3a.1 interaction (+ *P. infestans*): Although siRNA122 primarily targets mRNA from *StRES1*, it cleaves nucleotides encoding the residues tryptophan-321 and glycine-323, which are critical for the active interface between StRES1 (sensor) and StNRC3a.1 (helper) proteins. This cleavage disrupts the structural integrity of the sensor-helper complex, thereby weakening the immune response. In the absence of siRNA122 (exemplified by the transient CRISPR-mediated deletion), the StRES1-StNRC3a.1 complex remains intact, resulting in a restored resistance phenotype. Green dotted arrows indicate a resistant phenotype, whereas blue dotted arrows indicate susceptibility.

## MATERIALS AND METHODS

### StAGO15 cloning, constructs and plant transformation

Total RNA was isolated from potato using the RNeasy Plant Mini Kit (Qiagen), and cDNA was synthesized using the Maxima cDNA Synthesis Kit (Thermo Fisher Scientific). The *StAGO15* (PGSC0003DMT400039204) CDS was cloned from potato cv. Desirée and fused at the 3′ end to the *GFP* sequence in the binary plasmid pGWB505 harboring the 35S promoter (Nakagawa et al., 2007) and then transformed into the *Agrobacterium* strain GV3101 containing pSoup (Hellens *et al.,* 2000). RNAi primers for producing *StAGO15*-knockdown potato lines were designed using WMD3 (Web MicroRNA Designer http://wmd3.weigelworld.org/cgi-bin/webapp.cgi). The RNAi vector pRS300 (Addgene) and the *StAGO15* amiRNA precursor were ligated with the binary plasmid pGWB406 (Nakagawa et al., 2007) and transformed into *Agrobacterium* strain C58 containing pSoup (Hellens et al., 2000). Transformation into potato cv. Desirée was performed as previously described (Jahan et al., 2015). All generated constructs were confirmed *via* Sanger sequencing (Eurofins Genomics). The primers used for cloning are listed in Table S7. All potato primers used were based on genome version 6.1 (Pham et al., 2020).

### RNA coimmunoprecipitation, sequencing, and data analysis

Leaves with lesion sizes between 8 and 12 mm from a potato line harboring *p35S:StAGO15-GFP* and inoculated with *P. infestans* (*Pi*) strain 3928A were collected 5 days post-inoculation (dpi). In parallel, leaf materials from water-inoculated Desirée potato were prepared. Leaf materials from at least three plants were used for each replicate. Three replicates for each sample comprising 2 g of pooled leaf material were used for coimmunoprecipitation according to the manufacturer’s protocol (Chromotek gtm-100, Planegg-Martinsried, Germany) with minor adjustments (Åsman et al., 2016). Six libraries were prepared at SciLifeLab (Solna, Sweden). Libraries were prepared using the Illumina TruSeq Small RNA Kit and sequenced on an Illumina NextSeq 2000 platform (P2, 2×50 bp) with a 51 nt Read1, 6 nt Index1, and 51 nt Read2 configuration. Bcl-to-FastQ conversion was performed using bcl2fastq_v2.20.0.422 from the CASAVA software suite with the Sanger/phred33/Illumina 1.8+ quality scale. Data analysis was performed as previously described (Persson Hodén et al., 2021), except that the Illumina R2 FASTA files were reversely complemented *via* fastx_reverse_complement in the FASTX-Toolkit (Hannon, 2010) and combined with R1 files before using PAREsnip2. The resulting R1 and R2 FASTA files were combined, followed by the prediction of cleavage sites using the smartPARE pipeline (Persson Hodén et al., 2021), applying the same optimized convolutional neural network (CNN) model (100 iterations, 100% validation accuracy, lowest validation loss of 0.10). Datasets of infected and control materials were compared to generate increased or decreased normalized fragment abundance (NFA) datasets (Dataset 1).

### Quantitative reverse transcription-polymerase chain reaction (qRT-PCR)

Transgenic plants *p35S:StAGO15-GFP* (OE), *p35S:amiStAGO15-GFP* (KD), and cv. Desirée (WT) were inoculated with *P. infestans* (strain 3928A) as described previously (Jahan et al., 2015). RNA was extracted from *P. infestans*-inoculated and water-inoculated control leaves.

cDNA was synthesized using a Maxima First Strand cDNA Synthesis Kit for RT-qPCR (Thermo Fisher Scientific). Maxima SYBR Green/Fluorescein Master Mix (Thermo Fisher Scientific) was used for qRT-PCR (iQ5 Thermal Cycle, Bio-Rad), with *StActin101* used as a reference gene (Liao et al., 2020). The primers used are listed in Table S7.

### Pathogens and inoculations

Potato plants (cv. Desirée) were inoculated with *Phytophthora infestans* strain 3928A as previously described (Jahan et al., 2015). *Alternaria solani* strain As112 (Sajeevan et al., 2023) was maintained on potato dextrose medium (PDA). The concentration of fungal conidia was adjusted to 25,000 conidia ml⁻¹ in sterile water. Leaf inoculation with 10 μl of the spore suspension was performed, and sterile water was used as control.

### Pathogen quantification of infected leaves

To quantify *P. infestans* DNA in infected leaves, quantitative polymerase chain reaction (qPCR) analysis was carried out essentially as described previously (Llorente *et al*., 2010). Genomic DNA from infected potato leaves was isolated using cetyltrimethyl ammonium bromide (CTAB) lysis buffer at 65 °C for 1 h, followed by phenol/chloroform extraction. For qPCR analysis, 20 ng of DNA was used, and at least three biological replicates were analyzed. *PiO8* was used as the reference gene. P. infestans DNA was quantified with *PiO8* primers as described by Llorente et al. (2010) and normalized to potato *StActin101*. *A. solani* DNA was quantified by qPCR with the *A. solani*-specific primers AsFor and AsRev, which target the internal transcribed spacer region and yield a 152 bp product (Leiminger et al., 2015). The primers used are listed in Table S7.

### 5′RNA ligase-mediated rapid amplification of cDNA ends (5′RLM-RACE)

5′RLM-RACE was performed as follows (McCue et al., 2013): GeneRacer RNA Oligo (Invitrogen) was ligated to the total RNA using T4 RNA ligase (Nordic Biolabs AB, Stockholm, Sweden). cDNA was synthesized, and polymerase chain reaction (PCR) amplification was performed. First, touchdown PCR with the GeneRacer 5′ primer and a gene-specific reverse primer was run. The GeneRacer 5′ nested primer was then used for a second PCR, where the first PCR products served as a template. After gel purification, the amplified products were cloned and inserted into pJET1.2/blunt (Thermo Fisher Scientific) and sequenced. The oligo sequences are listed in Table S7.

### Northern blot and hybridization

Total RNA was extracted from potato plant samples using TRIzol (Thermo Fisher Scientific) according to the manufacturer’s recommendations. Thirty micrograms of RNA enriched for the low-molecular-weight fraction (Kreuze et al., 2005) was resolved on denaturing 17% polyacrylamide gels and transferred to nylon membranes. Commercially synthesized antisense sRNA-specific oligonucleotides were labeled using a DIG oligonucleotide 3′-end labeling kit (Sigma‒Aldrich) following the manufacturer’s instructions. DIG-labeled 21 nt sRNAs and potato U6 snRNA probes were hybridized to the membranes at 38 °C, followed by washing and visualization (Tomassi et al., 2017). The probe primer sequences are listed in Table S7.

### Electrophoretic mobility shift assay (EMSA)

The coding sequences for *StAGO15*, *StAGO1a*, and *StAGO1b* were cloned and inserted into the pCold*-TF* vector (Takara Bio) and expressed in a His-tagged *E. coli* BL21 strain (Thermo Fisher Scientific). Expression was induced at 16 °C to increase yields, and proteins were subsequently purified using Pierce™ High-Capacity Ni-IMAC Magnetic Beads, EDTA compatible (Thermo Fisher Scientific). Control gels with different eluates were stained with Coomassie blue. Biotin-labeled small RNAs and unlabeled nucleotide probes were synthesized (GenCefe Biotech, Zhenjiang, China). In the binding assays, biotin-labeled siRNA probes were incubated with the recombinant proteins StAGO15, StAGO1a, and StAGO1b. Electrophoretic mobility shift assays using 4% polyacrylamide native gels were run according to the LightShift™ Chemiluminescent RNA EMSA Kit protocol (Thermo Fisher Scientific) to determine and analyze sRNA-protein binding affinity. All three proteins were tested; owing to their high molecular mass, StAGO1a and StAGO1b could not be resolved, and only StAGO15 could be assayed.

### Genomic editing and transient assay

For genome editing, *MITE* source data were extracted from Dataset 1 and Dataset 3, and the siRNA122 source region was targeted with paired sgRNAs (Table S7). sgRNAs were designed following Dölfors et al. (2024) using CRISPys (Hyams et al., 2018) and CRISPR MultiTargeter (Prykhozhij et al., 2015), with potential off-targets assessed using Cas-OFFinder (Bae et al., 2014). The dual-sgRNA cassette was cloned into pHSE401 (Xing et al., 2014) to target a 201-nt region containing the siRNA122 source sequence within a *MITE* on chromosome 11 (9,393,316–9,393,516, + strand).

The transient expression system was adapted from Hoshikawa et al. (2019), combining the pBYR2HS geminiviral replicon (Yamamoto et al., 2018) with *Agrobacterium tumefaciens* expressing GABA transaminase (Nonaka et al., 2017, 2019). As these plasmids were unavailable, both were reconstructed separately. The p19 silencing suppressor was supplied as pGD-p19, while the tomato GABA transaminase gene (*SlGABA-t*) was cloned into pGWB406 (Nakagawa et al., 2007) for constitutive expression.

All plasmids were introduced into *A. tumefaciens* AGL1 (pTiBo542) by freeze-thaw transformation (Weigel & Glazebrook, 2006). Leaves from six-week-old potato plants were infiltrated as described by Zhang et al. (2020), with Silwet L-77 (0.01%) included at the recommended concentration. Plants were kept in darkness for 24 h at 21 °C before pathogen inoculation. Control (empty-vector) and genome-edited infiltrations received identical construct combinations.

### Statistical analyses

Beside the bioinformatics analyses of the sequencing information, a two-tailed Student’s *t*-test was employed for pairwise comparisons. Experiments examining the influence of two different variables on three or more samples were analyzed using two-way analysis of variance (ANOVA). Phenotypic observations were monitored for two to three generations for consistency.

## Supporting information

Dataset 1. StAGO15-mediated small RNAs and their targets.

Dataset 2. Unpaired energy (UPE) and expectation scores of RACE candidates.

Dataset 3. Predicted siRNAs in MITEs.

## AUTHOR CONTRIBUTIONS

SS made the KD plants, analyzed sRNA targets, performed the 5’RACE, northern blot, transient potato assay with CRISPR/Cas9, and ran the Co-IP together with KPH. XH performed the 5’RACE, RT-qPCR, EMSA, and construct preparations. KPH performed all bioinformatics and made Co-IP together with SS. ZL generated the transgenic *StAGO15* OE plants. SS, XH, CD designed the experiments. CD, SS, KPH, and XH wrote the manuscript.

## ACKNOWLEDGMENTS

We would like to thank Dr. S. Bhattacharjee for support with the protein analysis. This work was supported by the Knut and Alice Wallenberg Foundation (KAW 2019.0062), the Swedish Research Council Formas, and the Swedish University of Agricultural Sciences.

## DATA AVAILABILITY STATEMENT

The material that supports the findings of this study are available on request from the corresponding author. The sRNA sequencing data are available through the National Center for Biotechnology Information (NCBI) Gene Expression Omnibus GEO submission: https://www.ncbi.nlm.nih.gov/geo/query/acc.cgi?acc=GSE244421

