## SUPPORTING INFORMATION for "Argonaute 15 regulates immunity of potato to filamentous pathogens through transposon-derived small RNAs"

**Figure S1.** Precursor and target site summary of the sRNA targets in potato.
**Figure S2.** Predicted sRNA targets in potato according to annotated gene functional categories in the potato genome.
**Figure S3.** Target transcripts after filtration with smartPARE.
**Figure S4.** *Alternaria solani phenotypes and pathogen DNA abundance of StAGO15 transgenic lines***.**
**Figure S5.** Relative expression of *MITE/SINE*-generated sRNAs of target mRNAs upon *P. infestans* inoculation.
**Figure S6.** Electrophoretic mobility shift assay of StAGO15.
**Figure S7.** Cleavage efficiency identified by 5' RACE.
**Figure S8.** *StNRC3a.1* transcription in transgenic *StAGO15* lines.
**Figure S9.** Protein docking analysis between StRES1 and StNRC3a.1.
**Figure S10.** *P. infestans* DNA content in *StAGO15* native-promoter lines.

**Table S1.** Number of reads generated through Illumina sequencing.
**Table S2.** Detected miRNAs in the pulldown materials listed in Table S1.
**Table S3.** Distribution of siRNAs from TE categories associated with StAGO15.
**Table S4.** sRNA nucleotides (%) and lengths generated by the TEs listed in Table S3.
**Table S5.** Details of *MITE/SINE*-generated sRNA targets in potato used for qPCR validation.
**Table S6.** Protein domains of siRNAs targeting *R* gene sequences.
**Table S7.** List of oligos used in different assays in this study.

**Dataset 1.** *StAGO15*-mediated small RNAs and their targets.
**Dataset 2.** Unpaired energy (UPE) and expectation scores of RACE candidates.
**Dataset 3.** Predicted siRNAs in *MITE*s.

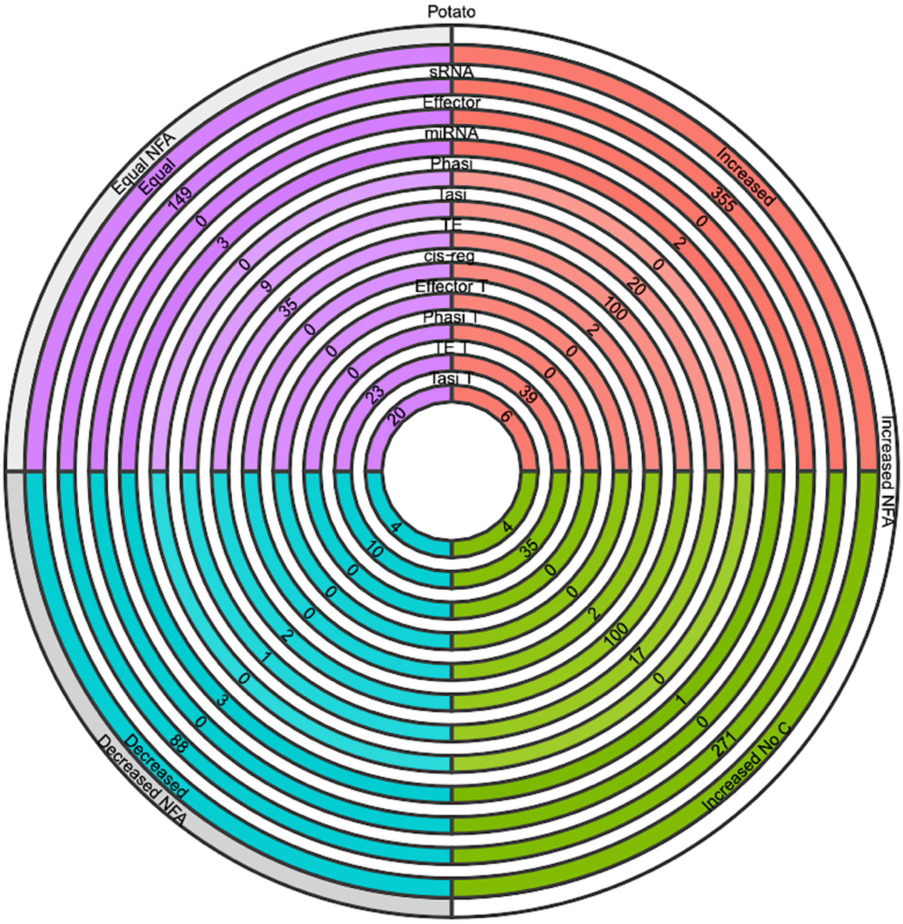


**Figure S1. Predicted precursor and small RNA targets in potato.** The number of precursor genes and targeted mRNAs is excluded. The gray ring specifies the normalized fragment abundance (NFA) level upon infection. Increased No C: represents the number of targets that are present in the infected material but not in the control samples. Increased: includes all the increased sites (including data from the "increased No C dataset. Subsequent rings toward the center correspond to different materials, RNA classes, and numbers of siRNAs related to each class. All classes denoted with a "T" at the end indicate that the class corresponds to the target site, whereas classes without "T" denote the precursor site. TE (transposons and repeats), cis-reg (cis-regulatory siRNAs), phasi (phasiRNAs), and tasi (tasiRNAs) are classified according to Axtell (2013).

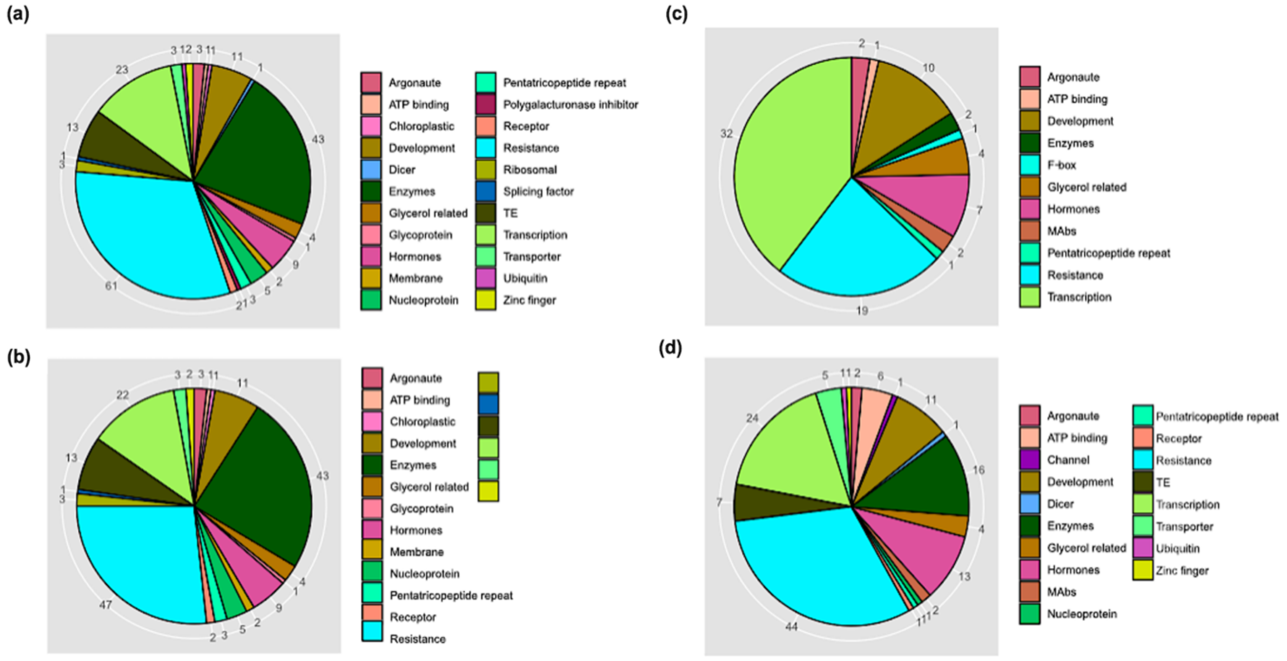


**Figure S2. Functional categories of genes targeted by predicted siRNAs in potato during *P. infestans* infection.** Predicted siRNA-targeted mRNAs in potato with different normalized fragment abundance (NFA) levels upon *P. infestans* infection in comparison with noninfected controls, organized according to annotated gene functional categories in the potato genome. (a) Increased NFA levels (more siRNAs). Thirty genes of unknown function and 14 genes not fitting into any of these categories were excluded. (b) Increased NFA, not present in the control (increased No C). Twenty-eight genes of unknown function and 14 genes not fitting into any of these categories were excluded. (c) Decreased NFA. Fifteen genes of unknown function and one gene not fitting into any of these categories were excluded. (d) Equal NFA levels. Thirty genes of unknown function and 12 genes not fitting into any of these categories were excluded.

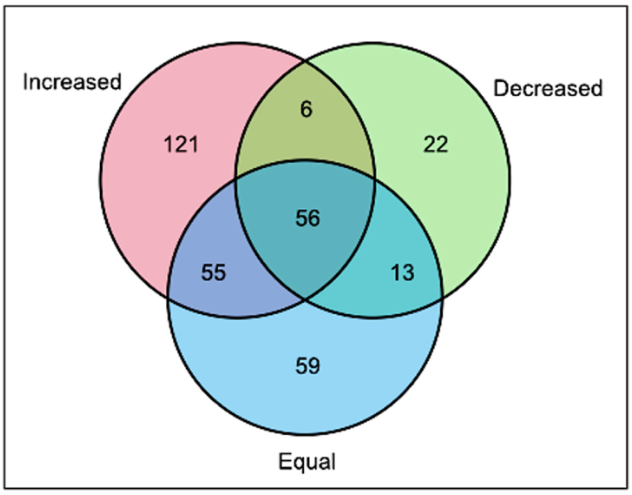


**Figure S3. Number of "true" target transcripts after data filtration with the smartPARE package** (Persson Hodén et al., 2021). Targeted transcripts in potato identified by siRNA with different normalized fragment abundance (NFA) levels upon *P. infestans* infection compared with the controls. The number of shared cumulative genes is indicated.

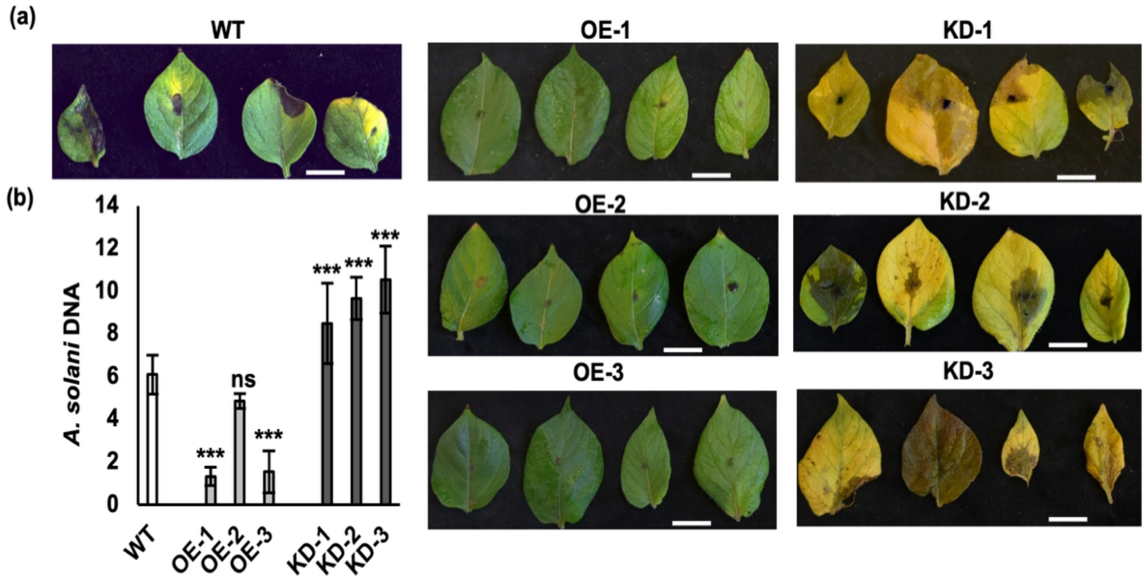


**Figure S4.** *Alternaria solani* phenotypes and pathogen DNA abundance of *StAGO15* transgenic lines. (**a**) Disease phenotypes of the wildtype (WT; cv. Desirée), three transgenic overexpression lines carrying p35S:StAGO15-GFP (OE-1, OE-2, OE-3) and three transgenic knockdown lines carrying p35S:amiStAGO15-GFP (KD-1, KD-2, KD-3) at 7 days post inoculation with *A. solani*. Four representative leaves are shown per genotype. Scale bars = 10 mm. (**b**) Relative *A. solani* DNA content in leaves of the same lines, quantified by qPCR with the *A. solani*-specific primers *AsFor* and *AsRev* (Leiminger et al., 2015) and normalized to potato *StActin101*. White bar, wildtype; light grey bars, overexpression lines; dark grey bars, knockdown lines. Error bars indicate the mean ± s.d. (n = 3). Asterisks indicate statistically significant differences from the wildtype according to two-tailed Student's t tests (*$*P < 0.05$, $**P < 0.01$, $***P < 0.001$*); ns, not significant.

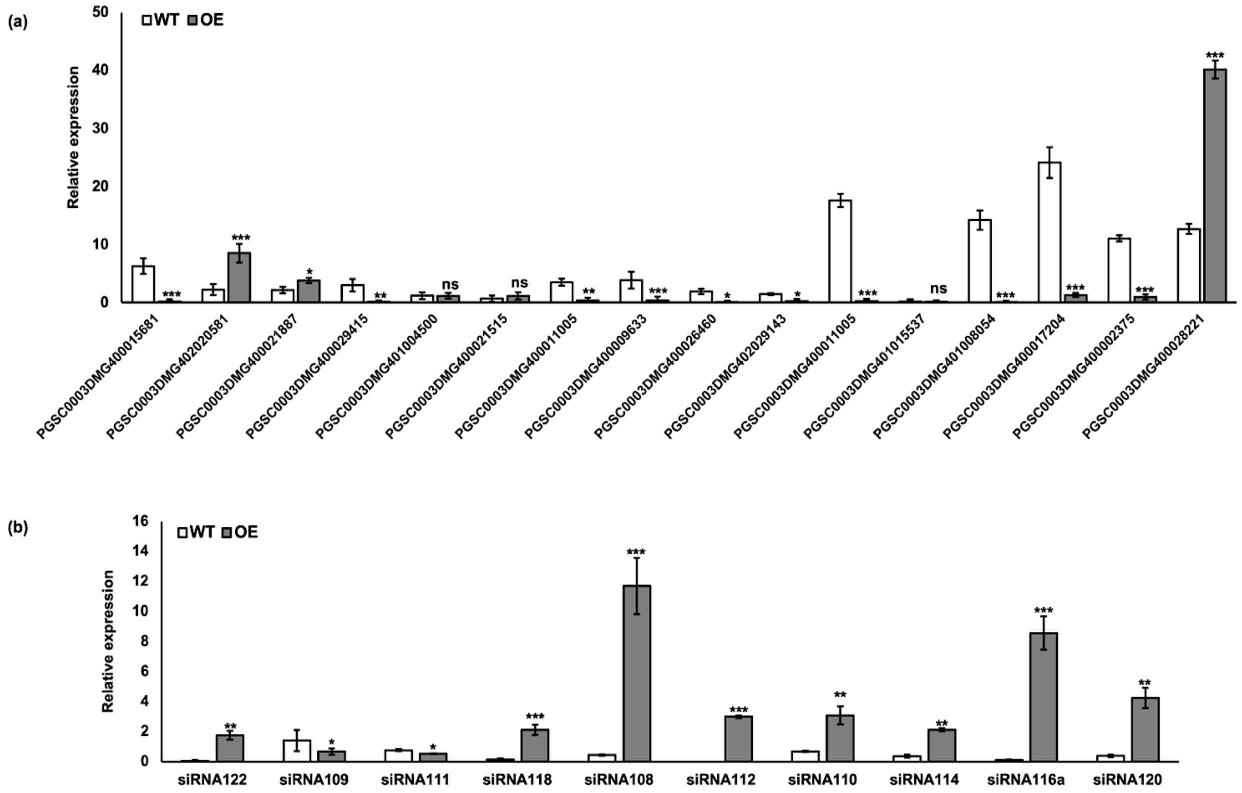


**Figure S5. Relative expression of *MITE/SINE*-generated sRNAs of target mRNAs upon *P. infestans* infection**. (a) Relative expression of target mRNAs in the wild-type (WT; cv. Desirée) (white bars) and transgenic overexpression lines (gray bars) inoculated with *P. infestans*. *StActi*n101 was used as a reference gene. (b) Relative expression of sRNAs in the wild-type (WT; cv. Desirée) (white bars) and transgenic overexpression lines (gray bars). sRNAs are listed in Table S5. *StActin101* was used as a reference gene. The error bars indicate the mean ± s.d. (n = 3). Asterisks indicate statistically significant differences for each line relative to WT at *$*P < 0.05$, $**P < 0.01$* and *$***P < 0.001$* according to two-tailed Student's t tests.

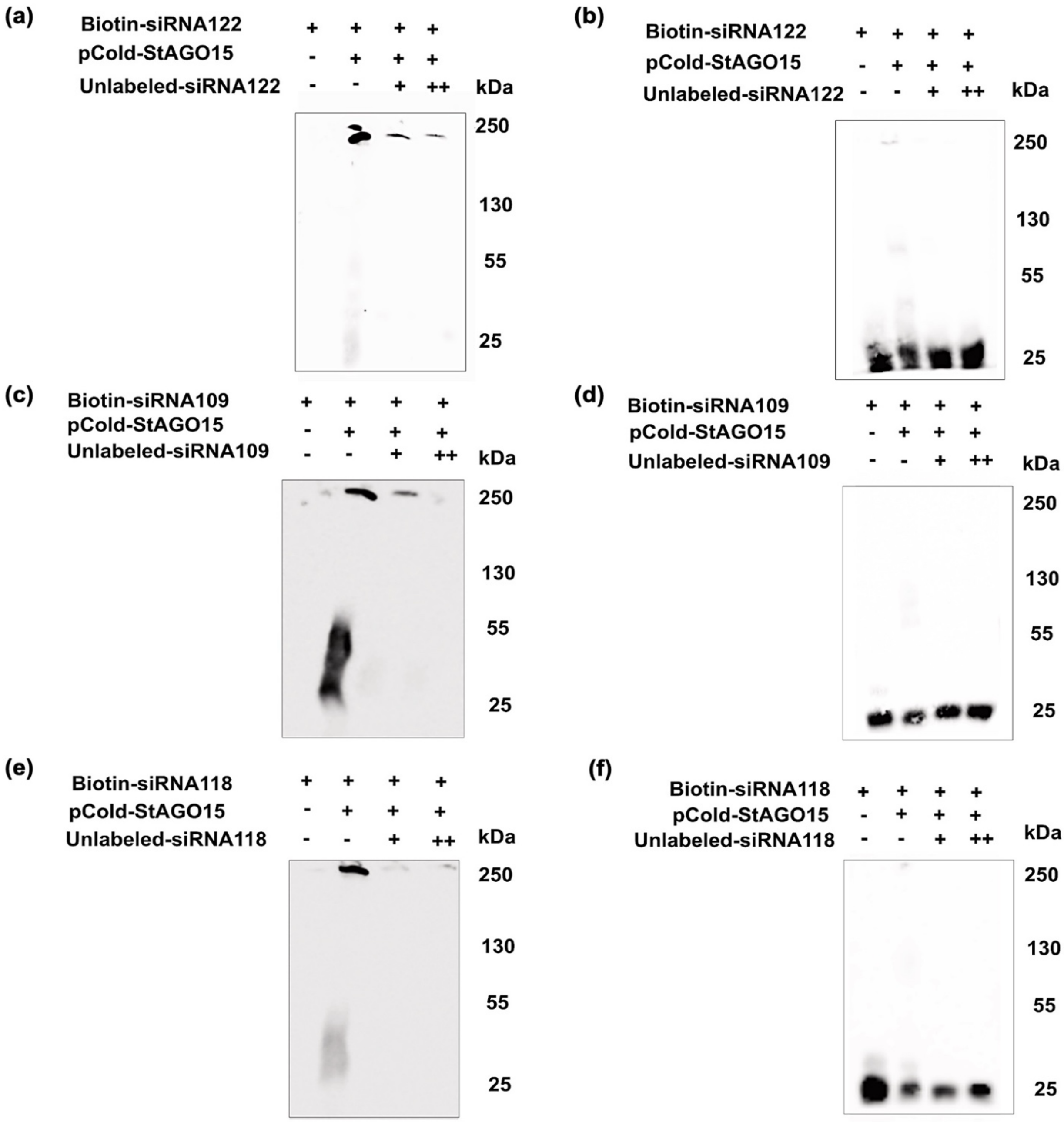


**Figure S6. Electrophoretic mobility shift assay (EMSA) analysis of StAGO15. EMSAs showing binding** between StAGO15 (approx. 158 kDa) and the (a) 21 nt siRNA122 (c) 22 nt siRNA109 and (e) 22 nt siRNA118 probes, with detection after 36 minutes, and the free (b) siRNA122, (d) siRNA109 and (f) siRNA118 probes after 12 minutes.

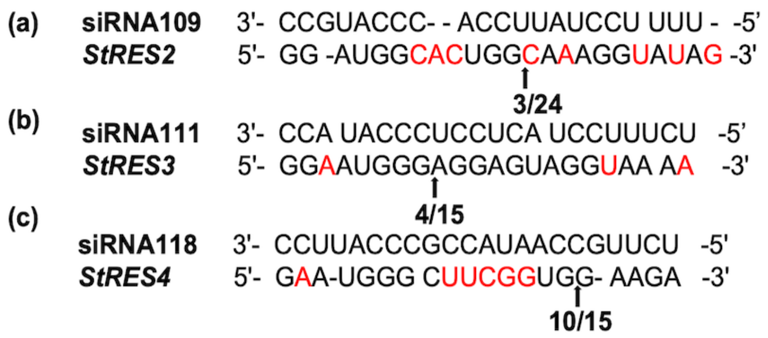


**Figure S7. Cleavage efficiency identified by 5' RACE.** Schematic drawing showing the alignment of siRNA109, siRNA111, siRNA118 with PGSC0003DMG402020581 (*StRES2*), PGSC0003DMG400021887 (*StRES3*) and PGSC0003DMG400029415 (*StRES4*), respectively, (TIR-NB-LRR category) at the predicted cleavage site during infection by *Phytophthora infestans*. Red nucleotide divergence sequence information between cv. *Desirée* and the reference genome. The arrow indicates the cleavage site identified.

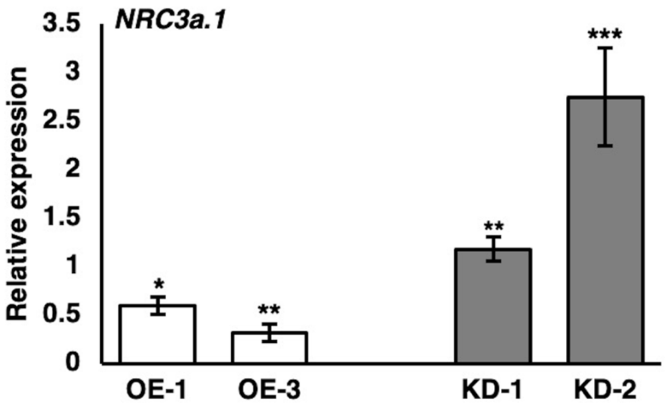


**Figure S8. *StNRC3a.1* transcription in transgenic *StAGO15* lines.** Relative expression of *StNRC3a.1* (Sotub05g007670.1) in transgenic overexpression lines (OE-white bars) and knockdown lines (KD, gray bars) compared with that in wild-type plants when inoculated with *P. infestans*. *StActin*101 was used as a reference gene. The error bars indicate the mean ± s.d. (n = 3). Asterisks indicate statistically significant differences $*P < 0.05$, $**P < 0.01$ and $***P < 0.001$ via two-tailed Student's t test.

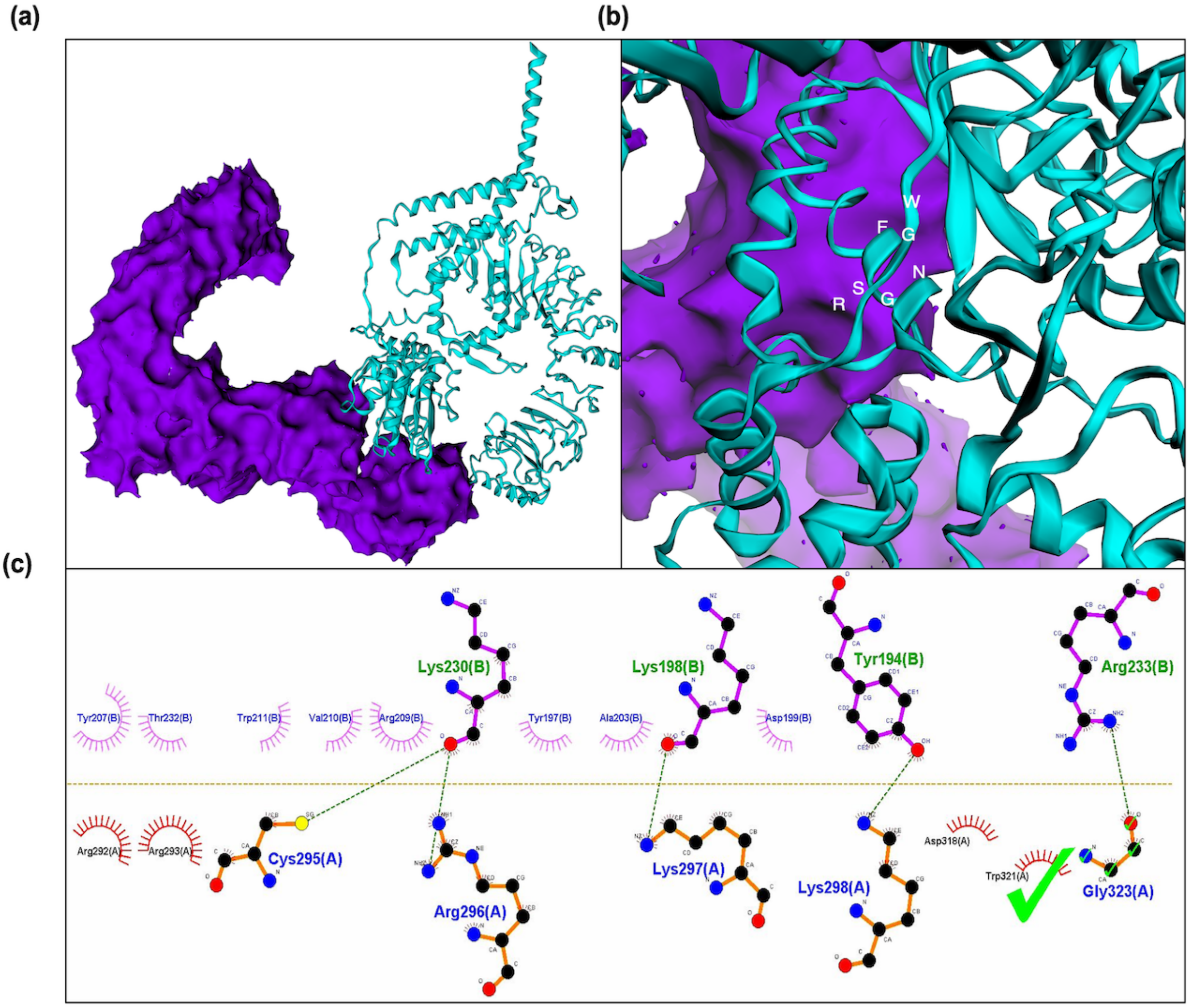


**Figure S9. Protein docking analysis between StRES1 and StNRC3a.1.** (**a**) Molecular docking visualization of StRES1 (violet) and StNRC3a.1 (cyan) via the EzMol interface (Reynolds et al., 2018). The docking results yield a HADDOCK score of 152.7 ± 19.6, with a cluster size of 30 and an RMSD of 6.5 ± 2.1 (Honorato et al., 2021, 2024). The key energy contributions include van der Waals (-52.4 ± 4.5), electrostatic (-176.0 ± 34.4), and desolvation (-0.7 ± 2.5) energies, with a high restraint violation energy (2409.2 ± 149.8) and a buried surface area of 2002.6 ± 132.1. The Z score of -0.5 places this cluster within a favorable range. (**b**) Enlarged view of (a) showing the RNA complementary site (highlighted in white residues), with key residues from chain A, including Trp321 and Gly323. (c) 2D LIG plot of the active interaction site via LigPlot+ (Laskowski & Swindells, 2011), highlighting Gly323(A) and Trp321(A) with a green tick mark. The plot illustrates critical residues from both chains, detailing hydrogen bonds and hydrophobic interactions.

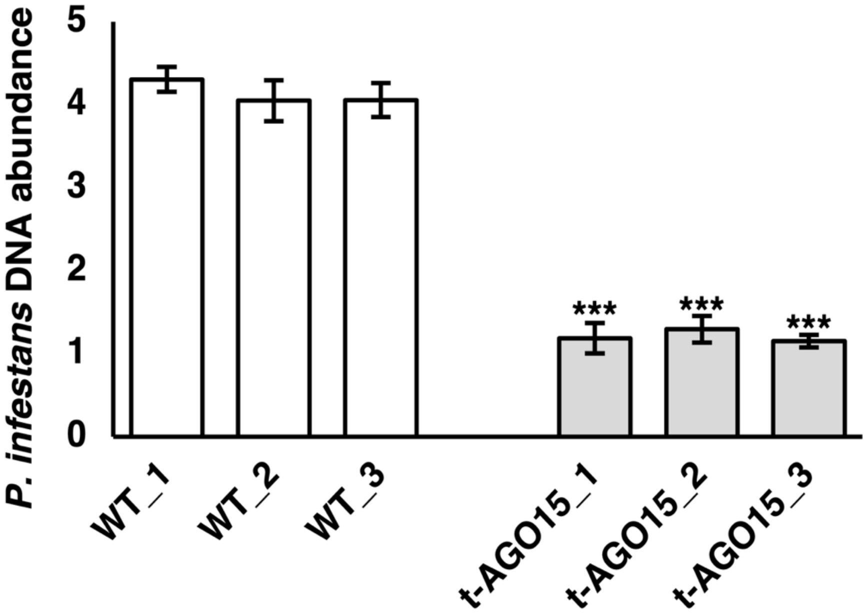


**Figure S10.** Relative *P. infestans* DNA content in leaves of three independent transgenic lines expressing *StAGO15* from its native promoter (t-AGO15_1 to t-AGO15_3) and of wildtype cv. *Desirée* (WT_1 to WT_3) at 5 dpi. *P. infestans* DNA was quantified with *PiO8* primers and normalized to potato *StActin101*. Error bars indicate the mean ± s.d. (n = 3). Asterisks indicate statistically significant differences from the wildtype *(***P < 0.001*) according to two-tailed Student's t tests.

**Table S1**. Number of reads after fusion of the Illumina libraries made after coimmunoprecipitation of leaves from a *p35:StAGO15-GFP* potato line $H_2O$ treated or inoculated by *P. infestans*. (Reverse complement was performed on the R2 libraries).

| **StAGO15 - Potato** | |
|---|---|
| Sample | No. of reads |
| C_1 | 3 687 202 |
| C_2 | 5 398 362 |
| C_3 | 4 117 395 |
| I_1 | 5 303 893 |
| I_2 | 2 498 668 |
| I_3 | 3 635 328 |
| **StAGO15 – *P. infestans*** | |
| Sample | Reads |
| I_1 | 1 386 411 |
| I_2 | 375 046 |
| I_3 | 501 507 |

I = reads from the *P. infestans* inoculated ~~*p35S:StAGO15-GFP*~~ leaves.

C = reads from the $H_2O$ inoculated ~~*p35S:StAGO15-GFP*~~ leaves.

**Table S2.** Detected potato miRNAs in the pulldown materials listed in Table S1.

| **miRNA** | **Closest homolog in miRBase[1]** | **$H_2O$** | ***P. infestans*** | **p-value[2]** |
|---|---|---|---|---|
| UUGACAGAAGAUAGAGAGCAC | stu-miR156c | 0 | 2 | 0.19 |
| AAGCUCAGGAGGGAUAGCACC | stu-miR390-5p | 1 | 0 | 0.42 |
| AUCAUGCGAUCUCUUCGGAAU | stu-miR393-3p | 0 | 1 | 0.42 |
| CGCUAUCCAUCUUGAGUUUUU | stu-miR390-3p | 1 | 0 | 0.42 |
| CUAGAUUCACGCACAAACUCG | vvi-miR403e | 1 | 0 | 0.42 |
| CUCACUUCUCUUUCUGUCAAUC | stu-miR156f-3p | 1 | 0 | 0.42 |
| GCACCGUCGCCGUCGGAG | hvt-miR-H4-3p | 1 | 0 | 0.42 |
| GUUGGCAAGUGGUCCUUGGC | ptc-miR169af | 1 | 0 | 0.42 |
| GUUUAUUAGCUAGAACUCC | osa-miR5157a-5p | 1 | 0 | 0.42 |
| UACCAACAAUUGAGAUAACAUC | stu-miR6025 | 1 | 0 | 0.42 |
| UCUUGCCAAUACCGCCCAUUCC | stu-miR482e-3p | 0 | 1 | 0.42 |
| UGACUAUCGAACCCUCACCAA | sly-miR1918 | 1 | 0 | 0.42 |
| UGUCGCAGAUGACUUUCGCCC | stu-miR1919-5p | 1 | 0 | 0.42 |
| UUCAACUUGCUCACUUGGAG | stu-miR8038a-3p | 2 | 0 | 0.42 |
| UUGCAACACACCUGAAUCGUC | stu-miR6149-5p | 0 | 1 | 0.42 |
| UUUGGAUUGAAGGGAGCUCUA | ath-miR159a | 1 | 0 | 0.42 |
| UCGGACCAGGCUUCAUUCCCC | stu-miR166d-3p | 1 | 1 | 0.47 |
| UUCCACAGCUUUCUUGAACUG | aau-miR396 | 3 | 3 | 0.48 |
| UGUGUUCUCAGGUCGCCCCUG | rco-miR398b | 3 | 3 | 0.51 |
| UGCACUGCCUCUUCCCUGGCU | stu-miR408b-3p | 1 | 1 | 0.63 |
| UAUGUUCUCAGGUCGCCCCUG | stu-miR398a-3p | 1 | 4 | 0.67 |
| UCGAUAAACCUCUGCAUCCAG | stu-miR162a-3p | 5 | 1 | 0.8 |
| UGCACAGCCUCUUCCCUGGUU | stu-miR408a-3p | 1 | 1 | 0.9 |
| UAUUGGCCUGGUUCACUCAGA | stu-miR171c-5p | 1 | 2 | 0.92 |

Abbreviations: Stu: *Solanum tuberosum*, sly: *Solanum lycopersicum*, ath: *Arabidopsis thaliana*, vvi: *Vitis vinifera*, rco: *Ricinus communis*, aau: *Acacia auriculiformis,* osa: *Oryza sativa*. hvt: Herpesvirus of turkeys, Ptc *Populus trichocarpa*.

1 = miRNA accession with highest resemblance according to Smith and Waterman (1981) in the miRBase database to the major miRNAs. Data retrieved from miRBase database September 2022.

2 = p-value of Student's t-test between the number of miRNAs in the pathogen infected vs. $H_2O$ samples.

**Table S3**. Distribution of siRNAs from TEs associated with *St*AGO15.

| **TE category** | **siRNA frequency based on TEs associated with *St*-AGO15 (%)** |
|---|---|
| *MITE* | 79.7 |
| *SINE* | 9.1 |
| *LTR* | 4.2 |
| *LTR / MITE* | 2.2 |
| *DNA / MITE* | 1.3 |
| *DNA/TcMar / MITE* | 1.2 |
| *LTR/Copia/LTR* | 0.7 |
| *LTR/Copia* | 0.5 |
| *LTR /SINE* | 0.5 |
| *SINE /MITE* | 0.4 |
| *DNA/hAT /MITE* | 0.2 |
| *LTR/Gypsy* | 0.2 |

**Table S4.** sRNA nucleotides (%) and lengths generated by the TEs listed in Table S3.

| **nt length** / **TE category** | 19 | 20 | 21 | 22 | 23 | 24 |
|---|---|---|---|---|---|---|
| *MITE* | 5.2 | 3.4 | 6.7 | 79.5 | 3.9 | 1.4 |
| *SINE* | 12.5 | 0 | 34.6 | 37.3 | 6.6 | 8.9 |
| *LTR* | 19 | 9.7 | 38.1 | 5.9 | 19.7 | 7.6 |
| *LTR \| MITE* | 7.4 | 0 | 0 | 76.6 | 16.1 | 0 |
| *DNA \| MITE* | 21.6 | 23.9 | 36.4 | 0 | 18.2 | 0 |
| *DNA/TcMar \| MITE* | 0 | 0 | 100 | 0 | 0 | 0 |
| *LTR/Copia \| LTR* | 0 | 0 | 45.8 | 31.2 | 22.9 | 0 |
| *LTR/Copia* | 100 | 0 | 0 | 0 | 0 | 0 |
| *LTR \| SINE* | 0 | 0 | 0 | 0 | 100 | 0 |
| *SINE \| MITE* | 0 | 0 | 0 | 0 | 100 | 0 |
| *DNA/hAT \| MITE* | 0 | 0 | 0 | 0 | 0 | 100 |
| *LTR/Gypsy* | 100 | 0 | 0 | 0 | 0 | 0 |

**Table S5.** Details of *MITE/SINE*-generated sRNA targets in potato used for qPCR validation.

| siRNA | Length | TE origin | Targets/Gene ID | Function | Tomato homologs[2] |
|---|---|---|---|---|---|
| **siRNA122** | | | | | Solyc11g069470.1.1/ |
| TCTGCTGCCATTACCAAACCA | 21nt | *MITE* | PGSC0003DMG400015681 | ***StRES1*[1]: Resistance gene** | 99.3% |
| **siRNA109** | 22nt | *MITE* | PGSC0003DMG402020581 | ***StRES2*[1]**: NBS-coding resistance gene | Solyc09g092410.2.1/ |
| TTTTTCCTATTCCACCCATGCC | | | | | 96.6% |
| **siRNA111** | 22nt | *MITE* | PGSC0003DMG400021887 | ***StRES3*[1]**: Bacterial spot disease resistance protein 4 | Solyc11g011060.2.1/ |
| TCTTTCCTACTCCTCCCATACC | | | | | 79.2% |
| **siRNA118** | 22nt | *SINE* | PGSC0003DMG400029415 | ***StRES4*[1]**: Nematode resistance protein | Solyc07g052770.2.1/ |
| TCTTGCCAATACCGCCCATTCC | | | | | 76.4% |
| **siRNA119** | 19nt | *LTR/ Copia* | PGSC0003DMG401004500 | Glycosyltransferase 2 | Solyc02g063000.2.1/ |
| ATCCACAATGTAGACATTT | | | | | 94.3% |
| **siRNA123** | 19nt | *LTR* | PGSC0003DMG400021515 | t-SNAREs | Solyc04g078130.2.1/ |
| AACAATAACATATCTAGTA | | | | | 96.4% |
| **siRNA108** | | | | | Solyc10g084770.1.1/ |
| CAGGGTCTATCAGAAACAACC | 21nt | *SINE* | PGSC0003DMG400011005 | Protein kinase | 99.5% |
| **siRNA114** | | | | | Solyc11g011350.1.1/ |
| TCTGCTGTCATTACCAAACCA | 21nt | N/A | PGSC0003DMG400009633 | N protein | 80% |
| **siRNA116a** | | | | | Solyc12g044410.1.1/ |
| TTCGGACCAGGCTTCATTCCC | 21nt | N/A | PGSC0003DMG400026460 | DNA binding protein | 99.6% |
| **siRNA116b** | | | | | Solyc08g066500.3.1/ |
| TTCGGACCAGGCTTCATTCCC | 21nt | N/A | PGSC0003DMG402029143 | Class III HD-Zip protein 8 | 99.8% |
| **siRNA120** | | | | | Solyc10g084770.1.1/ |
| AGCCAAGGGTCTATCGGAAAC | 21nt | *SINE* | PGSC0003DMG400011005 | Protein kinase | 99.5% |
| **siRNA125** | | | | | Solyc11g073130.1.1/ |
| CTCGGACTCTTCAAAAATGTC | 21nt | *MITE* | PGSC0003DMG401015537 | Tyrosine phosphatase | 97.7% |
| **siRNA112** | | | | | Solyc11g069470.1.1/ |

| | | | | | |
|---|---|---|---|---|---|
| TTCGGACCAGGCTTC ATTCCC | 21nt | N/A | PGSC0003DMG40 1008054 | DNA binding protein | 99.3% |
| **siRNA110** | | | | | Solyc09g090470.2.1/ |
| TGTGTCAGATCCTCC AAAAATA | 22nt | *MITE* | PGSC0003DMG40 0017204 | Oligopeptide transporter | 99.3% |
| **siRNA124** | | | | | Solyc09g061680.2.1/ |
| AAATTTTGTGTCGAG TCAAACT | 22nt | *MITE* | PGSC0003DMG40 0002375 | Pantoate:beta-alanine ligase | 96.8% |
| **siRNA126** | | | | | Solyc10g081560.2.1/ |
| CTTGAGAAATATAG GTAGGGCCG | 23nt | *MITE* | PGSC0003DMG40 0028221 | Tropinone reductase I | 98.1% |

[1] *StRES1-StRES4* = Selected for 5′ RACE validation of *StRES2*, *StRES3* and *StRES4* carry different top-hit annotations (NBS-coding resistance gene, Bacterial spot disease resistance protein 4, and Nematode resistance protein, respectively), belonging to the TIR-NB-LRR class referenced in Figure S7.

[2] Sequence similarity (%) with *S. tuberosum.*

**Table S6.** Protein domains of siRNAs targeting *R* gene sequences.

| Small RNA | Domains |
|---|---|
| siRNA122 | NB-ARC/ P-loop containing nucleoside triphosphate hydrolase |
| siRNA109 | NB-ARC |
| siRNA111 | P-loop containing nucleoside triphosphate hydrolase |
| siRNA118 | NB-ARC/ P-loop containing nucleoside triphosphate hydrolase |

**Table S7**. List of oligos used in different assays in this study.

| Oligo name | Sequence 5' to 3' |
|---|---|
| **Cloning: Gene knockdown** | |
| Ami-StAgo15-OligoA | GGGGACAAGTTTGTACAAAAAAGCAGGCTTTCTGCAAGGCGATTAAGT TGGGTAAC |
| Ami-StAgo15-OligoB | GGGGACCACTTTGTACAAGAAAGCTGGGTCCTAGCGGATAACAATTTCA CACAGGAAACA |
| StAGO15_1_I miR-s | GATTGATTATTGGATCGCCTCAGTCTCTCTTTTGTATTCC |
| StAGO15_1_II miR-a | GACTGAGGCGATCCAATAATCAATCAAAGAGAATCAATGA |
| StAGO15_1_III miR*s | GACTAAGGCGATCCATTAATCATTCACAGGTCGTGATATG |
| StAGO15_1_IV miR*a | GAATGATTAATGGATCGCCTTAGTCTACATATATATTCCT |
| StAGO15_2_I miR-s | GATTCTATCGGCTTTTGGCGCGATCTCTCTTTTGTATTCC |

| | |
|---|---|
| StAGO15_2_II miR-a | GATCGCGCCAAAAGCCGATAGAATCAAAGAGAATCAATGA |
| StAGO15_2_III miR*s | CACCACCTCCCAAAGAAATTCC |
| StAGO15_2_IV miR*a | ATCTTCTTTCCCTTGACCCCC |
| **Cloning: Gene overexpression** | |
| StAGO15_attB_FP_N | GGGGACAAGTTTGTACAAAAAAGCAGGCTATGGCGTCTAAGGAGTTATTCCC |
| StAGO15_attB_RP_N | GGGGACCACTTTGTACAAGAAAGCTGGGTCCTATCAGCAGAAAAACATTGTATCGCG |
| StAGO15_pENTR11_NotI_N | GCGGCCGCGCAGAAAAACATTGTATCGCGCAC |
| StAGO15_pENTR11_XhoI_N | CTCGAGGCAGAAAAACATTGTATCGCGCAC |
| SlGABA-T2_attB_FP_N | GGGGACAAGTTTGTACAAAAAAGCAGGCTATGGCCAAGACTAATGG |
| SlGABA-T2_attB_RP_N | GGGGACCACTTTGTACAAGAAAGCTGGGTTTACTTCTTCTGAGACTTTAATTCTTCCACTC |
| **Gene expression** | |
| StAgo15_FP | GTCGAGCCAAAAGCCGATAGA |
| StAgo15_RP | TCCTTGTTTCCCACTCTCAACTG |
| StACT101_FP | GCCTCCTGAACGGAAGTACA |
| StACT101_RP | AATGGAAGGACCGGATTCAT |
| StAGO1a_FP | GTGCGGAAGAGGAGAACTGA |
| StAGO1a_RP | ACCATATCCTCCACGCTGAG |
| StNRC3a.1_FP | GATGCAATTACTAATCGACAACGC |
| StNRC3a.1_RP | GGATTTAGCAGCTTGTTTGAGGA |
| StFbox1_FP | GGCCGTGGGATGAATTGTTA |
| StFbox1_RP | GGTTTTCAGGGCAGCGATTT |
| **Target validation** | |
| Universal RP_Stem_Loop | CCA GTG CAG GGT CCG AGG T |
| PG53049_FP | TGATGAGCTGGCACAAGACA |
| PG53049_RP | CATGCATCACCAAATAACGGCA |
| PG44306_FP | GCACTCTACACCTTAGCCGT |
| PG44306_RP | AGTGACCAGCCCACATTGTC |
| PG56338_FP | TATGGCGTGGATGCAACAGA |
| PG56338_RP | GATTTGCGGCAGCAGTTAGG |
| PG20801_FP | GCTCAGCAGCATAGGGAGAG |
| PG20801_RP | CTCGCGACACCTTCTGTTCT |
| PG24927/29_FP | ACCAAGGCAGTGGAAGCAAT |
| PG24927/29_RP | TCTTGGCAATTGGAGTCGGG |
| PG68035/37/38_FP | GATCGCCCTTCCTGGTATCG |
| PG68035/37/38_RP | TTAACTGCCTCAGTTGGCGT |

| | |
|---|---|
| PG74934_FP | GCACCAGCTCGTGACTTTTG |
| PG74934_RP | GAACCACCTCCTTCACAGGG |
| PG75625_FP | GGTGGATTGAATGGCCGAAC |
| PG75625_RP | TCTTCGTTAGCCCAAGGCTC |
| PG11466_FP | TGATTGAGCTTGCATGGGGT |
| PG11466_RP | TGACACGTGTATCGTTCCCA |
| PG28589_FP | CACAGAGAATGGCTGGCTGA |
| PG28589_RP | GGGGCAGTGCAAACTGTAAA |
| PG28592_FP | GGAAGGCCATCTTACTGCCA |
| PG28592_RP | TCGTCCATTGTTGGCCTTGA |
| PG40528_FP | CTGCTGATGGATTGGGGGAG |
| PG40528_RP | TCACAGCATTCATGGCGTTT |
| PG55418_FP | GGTTGGTATGGTGCCGTTCA |
| PG55418_RP | CGTTAGCAAACTCACCAATCAGG |
| PG06113_FP | GAGTATGAGAGGCCAAGGCA |
| PG06113_RP | ACCCCACCAGGTACAGAGTT |
| PG40152_FP | GGGGTTTATGCTTCTGCTGGT |
| PG40152_RP | GAGTTCGTCCACTCCATTGC |
| PG72532_FP | CCAGCAGTCATTTTAACCCCACAA |
| PG72532_R | CTGTAAATCCACCATCGGCCC |
| **siRNA validation** | |
| siRNA_108_ST-RT | GTCGTATCCAGTGCAGGGTCCGAGGTATTCGCACTGGATACGACGTCCCA |
| siRNA_108_FP | GGTGCCAGGGTCTATCAGAAACAA |
| siRNA_112_ST-RT | GTCGTATCCAGTGCAGGGTCCGAGGTATTCGCACTGGATACGACAAGCCT |
| siRNA_112_FP | GGTGCTTCGGACCAGGCTTCATTC |
| siRNA_114_ST-RT | GTCGTATCCAGTGCAGGGTCCGAGGTATTCGCACTGGATACGACAGACGA |
| siRNA_114_FP | GGTGCTCTGCTGTCATTACCAAA |
| siRNA_116a_ST-RT | GTCGTATCCAGTGCAGGGTCCGAGGTATTCGCACTGGATACGACAAGCCT |
| siRNA_116a_FP | GGTGCTTCGGACCAGGCTTCATTC |
| siRNA _118_ST-RT | GTCGTATCCAGTGCAGGGTCCGAGGTATTCGCACTGGATACGACAGAACG |
| siRNA_118_FP | GGTGCTCTTGCCAATACCGCCCAT |
| siRNA _120_ST-RT | GTCGTATCCAGTGCAGGGTCCGAGGTATTCGCACTGGATACGACTCGGTT |
| siRNA_120_FP | GGTGCAGCCAAGGGTCTATCGGAA |
| siRNA _122_ST-RT | GTCGTATCCAGTGCAGGGTCCGAGGTATTCGCACTGGATACGACAGACG |
| siRNA_122_FP | GGTGCTCTGCTGCCATTACCAAAC |

| | |
|---|---|
| siRNA _109_ST-RT | GTCGTATCCAGTGCAGGGTCCGAGGTATTCGCACTGGATACGACGGCAT G |
| siRNA_109_FP | GCGGCGGTTTTTCCTATTCCACC |
| siRNA _111_ST-RT | GTCGTATCCAGTGCAGGGTCCGAGGTATTCGCACTGGATACGACGGTAT G |
| siRNA_111_FP | GCGGCGGTCTTTCCTACTCCTCC |
| siRNA _118_ST-RT1 | GTCGTATCCAGTGCAGGGTCCGAGGTATTCGCACTGGATACGACGGAAT G |
| siRNA_118_FP1 | GCGGCGGTCTTGCCAATACCGCC |
| **RACE** | |
| GbRacr5 Primer | CGACTGGAGCACGAGGACACTGA |
| GnRacr5NestdPrimer | GGACACTGACATGGACTGAAGGAGTA |
| PG28589_500bp-RP | GCTTGGCATTGTAGTCCTGTC |
| PG28589_300bp-RP | GATGATATTCACATACCGAGTCCAG |
| PG28592_500bp-RP | GGTACTTGGATTTCGCGGGG |
| PG28592_300bp-RP | TGCTGATTTACCTGCCGGAC |
| PG528_500bp-RP | GCTGAGTTCTTCCCTTTGCAC |
| PG528_300bp-RP | CTCACGAAAGGAAGAACCCCA |
| GeneRacerRNAOligo | CGACTGGAGCACGAGGACACTGACATGGACTGAAGGAGTAGAAA |
| siRNA111_300bp_RP | AACGAAGATCCCCTGCTAG |
| siRNA111_150bp_ RP | TTGCAGAGAGTACATTCTAT |
| siRNA109_250bp_ RP | TCAGAACTTTGCTCTTTTATGCC |
| siRNA109_150bp_ RP | GCCGGGAAAGATGTAACATTG |
| siRNA118_600bp_ RP | TCGATGAGGTAGCTTATTCCT |
| siRNA118_500bp_ RP | CAAGCATGCTATATCAAGGAATG |
| siRNA118_250bp_ RP | GCAAAACAGTTCAAGAGCTTCG |
| siRNA118_150bp_ RP | CTACTACCCCTGCCAAACCA |
| pJET1.2 FP Seq. | CGACTCACTATAGGGAGAGCGGC |
| pJET1.2 RP Seq. | AAGAACATCGATTTTCCATGGCAG |
| **Northern blot** | |
| 25 miRNA Ladder | AGAGCAGTGGCTGGTTGAGATTTAA |
| 22 miRNA Ladder | AGCAGTGGCTGGTTGAGATTTA |
| 17 miRNA Ladder | CAGTGGCTGGTTGAGAT |
| 21 miRNA Probe | AAATCTCAACCAGCCACTGCT |
| siRNA_122_528_Probe | TGGTTTGGTAATGGCAGCAGA |
| StU6_Probe | GGAACGATACAGAGAAGATTAGC |
| **EMSA** | |
| StAGO15_F | GGGGTACCATGGCGTCTAAGGAGTTATTC |
| StAGO15_R | CCCAAGCTTTCAGCAGAAAAACATTGTATCGCGCACA |
| StAGO1a_F | GGGGTACCATGGTGCGGAAGAGGAGAAC |
| StAGO1a_R | CCCAAGCTTCTAACAATAGAACATCACCCTTTTGAC |

| | |
|---|---|
| StAGO1b_F | GGGGTACCATGGTGAGGAAGAGGAGAACTG |
| StAGO1b_R | CCCAAGCTTCTAGCAATAGAACATGACCCTCT |
| siRNA122_Prob-3'biotin | TCTGCTGCCATTACCAAACCA |
| siRNA118_Prob-3'biotin | TCTTGCCAATACCGCCCATTCC |
| siRNA109_Prob-3'biotin | TTTTTCCTATTCCACCCATGCC |
| ***P. infestans*** **quantification** | |
| Pin_O8_FP | CAATTCGCCACCTTCTTCGA |
| Pin_O8_RP | GCCTTCCTGCCCTCAAGAAC |
| Pin_ACT_FP | CATCAAGGAGAAGCTGACGTACA |
| Pin_ACT_RP | GACGACTCGGCGGCAG |
| ***A.solani*** **quantification** | |
| A.solani_FP | TCCGTAGCTGAACCTGCGG |
| A.solani_RP | TGGGTTGGTCCTTGTGGTG |
| **CRISPR** | |
| siRNA122_FP | ATATATGGTCTCGATTCTGTCATTACCAAACCAACTGTTTTAGAGCTAG AAATAGC |
| siRNA122_RP | ATTATTGGTCTCGAAACAGTTGGTTTGGTAATGACAGAATCTCTTAGTC GACTCTAC |
| siRNA109_FP | ATATATGGTCTCGATTGATTTGGGAAGCTTTTGGAGGTTTTAGAGCTAG AAATAGC |
| siRNA109_RP | ATTATTGGTCTCGAAACTTGGCATGGGTGGAATAGGAAATCTCTTAGTC GACTCTAC |
| siRNA111_FP | ATATATGGTCTCGATTGGAAGTTGTTGGTTGTGGGTGTTTTAGAGCTAG AAATAGC |
| siRNA111_RP | ATTATTGGTCTCGAAACTTTTGATTTGGAAAAAGATAAATCTCTTAGTC GACTCTAC |
| siRNA118_FP | ATATATGGTCTCGATTGGAAGGGGATGAGTGTTTCGGTTTTAGAGCTAG AAATAGC |
| siRNA118_RP | ATTATTGGTCTCGAAAC CCCCGAAACACTCATCCCCTAATCTCTTAGTCGACTCTAC |
| U6-26p-FP Seq. | TGTCCCAGGATTAGAATGATTAGGC |


### ACKNOWLEDGEMENTS


We want to acknowledge the FP7 WeNMR (project# 261572), H2020 West-Life (project# 675858), the EOSC-hub (project# 777536) and the EGI-ACE (project# 101017567) European e-Infrastructure projects for the use of their web portals, which make use of the EGI infrastructure with the dedicated support of CESNET-MCC, INFN-LNL-2, NCG-INGRID-PT, TW-NCHC, CESGA, IFCA-LCG2, UA-BITP, TR-FC1-ULAKBIM, CSTCLOUD-EGI, IN2P3-CPPM, CIRMMP, SURFsara and

NIKHEF, and the additional support of the national GRID Initiatives of Belgium, France, Italy, Germany, the Netherlands, Poland, Portugal, Spain, UK, Taiwan and the US Open Science Grid.
